# Herbivorous fish of Sinai marine protected areas (Gulf of Aqaba): Structure baseline for potential thermal stress impact management

**DOI:** 10.1101/2021.09.17.460804

**Authors:** Khaled M. El-Haddad, Abdel-Hamid A. M. Ali, Mohammed S. Abdel-Rahman, Saad Z. Mohammed, Fagr Kh. Abdel-Gawad, Giulia Guerriero, Tarek A. Temraz

## Abstract

Herbivorous reef fishes are critically important for maintaining the health and resilience of coral reefs. In the Egyptian Gulf of Aqaba (GoA), many of these herbivores are important fishery targets, however, little is known about their biomass and size structure patterns. Here, we conduct a large-scale (∼250 km of coastline) assessment of the status of herbivorous fish populations on 30 reef sites from eight coastal regions in the GoA. These regions are subjected to three levels of fishing and protection, ranging from almost no fishing (No-take, NT), and moderately fished (via Gear-restriction, GR) to heavily fished (Open-access, OA). We found that the NT fishery reserve was the most effective in maintaining herbivorous fish richness, body-size, and biomass across all functional groups, though numerical density was not. Total herbivore biomass was 4.3 and 2.8 times higher on NT reefs and GR reefs respectively than on OA reefs. Among GoA regions, only Ras Mohammed and Sharm El-Sheikh (unfished), and Nabq (moderately fished) met the global mean herbivore biomass target of ∼30 kg/500 m^2^. Our work also highlights three important results regarding the impacts of fishing and fisheries management on herbivorous fish populations: (i) size structure was heavily skewed toward smaller individuals, with fishes less than (20 cm) accounting for 81.1% of the population in heavily fished OA reefs vs. 52.8% at unfished NT reefs, (ii) biomass of larger herbivore individuals (>35 cm) accounted for less than 1% of the total biomass in OA reefs vs. 37% at NT reefs, and (iii) large-bodied target species, *Cetoscarus bicolor, Naso unicorn*, and *Kyphosus* spp., accounting for 15.9% of the total biomass at NT reefs on average, while they were virtually absent from OA reefs. Collectively, these findings suggest that many principal fishery species in the central-northern regions of the GoA have been overexploited, which in turn can lead to reef degradation. Therefore, there is an urgent need to develop a participatory management approach and enforce restrictions on destructive fishing gear in order to promote recovery of herbivore biomass.

**Highlights:**

- Large-scale assessment of the status of herbivorous fishes in the northern Red Sea.
- Comparison of herbivorous fish assemblages across three levels of fishing pressure.
- Large-bodied herbivore species were either rare or absent at heavily fished reefs.
- Loss of these key herbivores may boost algal dominance, thus diminish reef functionality.
- Promoting local community participation in the fishery management process.

## 1. Introduction

Coral reef ecosystem are among the world’s most structurally complex, biodiverse and valuable ecosystems, supporting nearly a third of all marine fish species (Wilkinson, 2008; Graham and Nash, 2013; Fisher *et al*., 2015). However, this unique ecosystem is among the most vulnerable to local stressors mostly from overfishing (Burke *et al*., 2011). While more than 25% of the world’s coral reefs are located within marine protected areas (MPAs) boundaries, only 6% of reef areas are within adequately managed MPAs (Burke *et al*., 2011). Worldwide, many people depend on coral reefs as a source of food and income, both from reef fisheries and reef-based tourism activities (Wilkinson, 2008). Approximately 75% of the world’s reefs are found in developing countries and the rapid growth of the reef-based tourism sector has contributed to increasing demand for reef fishes, which, in turn, has significantly depleted many fish populations (Ashworth and Ormond, 2005; Burke *et al*., 2011). Indeed, managing the ever-increasing demand for coral reef fisheries while maintaining the functionality and ecological resilience of reefs is one of the most important challenging issues regarding corals’ present and future (Pratchett *et al*., 2014; Chung *et al*., 2019).

The GoA in the northern Red Sea is home to one of the world’s most diverse high-latitude reefs and a coral refuge from global stressors, yet they are also under increasing pressure from local stressors, in particular overfishing and destructive fishing methods (Burke *et al*., 2011; Fine *et al*., 2013; Osman *et al*., 2018). Egypt’s reefs have a national economic value due to their importance for local artisanal fisheries and tourism, ranking as one of the top ten countries with the highest total coral reef tourism value, accounting for $5.5 billion annually (Spalding *et al*., 2017). Coral reefs along the Egyptian GoA coast have been fished by the local Bedouin community for generations as part of their traditional activities (Poonian, 2020). To protect and conserve these unique reefs and coral reef fisheries, Egypt established a network of three connected MPAs in South Sinai, covering the entirety of Egypt’s GoA (Pearson and Shehata, 1998). However, their management effectiveness has diminished over time due to inadequate staffing and financial resources, especially in the past decade (Mabrouk, 2015; Samy-kamal, 2015). Additionally, non-compliance and weak enforcement of fisheries regulations in the MPAs may constitute a threat to conservation goals and hence the efficacy of the MPAs (Advani *et al*., 2015; Gill *et al*., 2017).

Fisheries management varies widely across GoA regions. In Ras Mohammed and Sharm el-Sheikh (south GoA), all forms of nearshore fishing activities are prohibited (Mabrouk, 2015). However, in Nabq (north of Sharm el-Sheikh), fishing is permitted for local Bedouin using their traditional gears (e.g., gillnets and trammel nets) that principally target herbivorous fish species entering the reef flat with the rising tides to feed on algae (Galal *et al*., 2002). Whereas, in Abu Galum and Dahab (central GoA), fishing is allowed in certain locations and prohibited at dive sites (Hasler and Ott, 2008). In contrast, fisheries management appear to be more permissive in Nuweiba and Taba (north GoA), where a variety of fishing gears are used (e.g., hook and line, nets, traps, spear guns) throughout these regions (Tilot *et al*., 2008; Poonian, 2020).

Local management of herbivorous fish populations (via size limits, gear restrictions, or spatial closures) has been widely acknowledged as an effective tool that reef managers can use to protect or restore healthy herbivore populations and indirectly maintain reef health and resilience (Bozec *et al*., 2016; Williams *et al*., 2016, 2019; Weijerman *et al*., 2018; Chung *et al*., 2019). Herbivorous fish species are largely classified into four main functional groups, namely (grazers, browsers, scrapers, and excavators), their ecological roles are critical in controlling coral-algal spatial competition (Burkepile and Hay, 2010, 2008; Cheal et al., 2010; Green and Bellwood, 2009). Therefore, a growing number of recent studies have recognized that monitoring and effective management of herbivorous fishes are important tools that reef managers can use for maintaining coral reef resilience. Specifically, it has been recommended that fish biomass be included in monitoring programs as the most useful indicator or response metric to evaluate the status of coral reef fisheries and set practical management targets (Green and Bellwood, 2009; Maynard *et al*., 2015; McClanahan *et al*., 2015, 2019, 2021; Chung *et al*., 2019; Williams *et al*., 2019; Campbell *et al*., 2020).

Herbivorous reef fishes have been studied on many coral reefs throughout the world (Cure et al., 2021; Helyer and Samhouri, 2017; Robinson et al., 2020; Shantz et al., 2020) and the Red Sea (Alwany *et al*., 2009; Afeworki *et al*., 2013; Khalil *et al*., 2013, 2017; Kattan *et al*., 2017). Whereas, a series of studies have described the status of herbivorous fishes in South Sinai MPAs (SSMPAs) in the GoA (Ashworth and Ormond, 2005; Tilot *et al*., 2008; Advani *et al*., 2015; Naumann *et al*., 2015; Reverter *et al*., 2020). Nevertheless, these studies were generally limited by the fact that they focused on a specific region or a fish group. Furthermore, quantitative information on the biomass and size structure of the functional groups of herbivorous reef fishes on the Egyptian GoA reefs is largely missing, hampering attempts to measure the effectiveness of SSMPAs in maintaining or enhancing herbivorous fish communities after several decades of its establishment. The present study, therefore, is intended to provide a better understanding whether SSMPAs and their fisheries management systems (FMSs) were effective in protecting the herbivorous fish richness, density, biomass, and size; that are essential to sustain the condition of coral reef ecosystems.

## 2. Methods

### 2.1. Study sites and survey design

Underwater visual censuses of herbivorous reef fishes were carried out between April-September 2017. A total of 30 reef sites from eight regions were surveyed. These sites were distributed along 250 km of coastline in the northern Red Sea and the GoA (Figure 1& Table S1). In GoA regions, three FMSs were categorized based on the prevalent human activity, level of conservation and protection, fishing intensity, and compliance with fishery regulations (Table 1). The southern GoA regions (Ras Mohammad and Sharm El-Sheikh) are classified as NT, where near-shore fishing is prohibited, the central GoA regions (Nabq, Dahab and Abu Galum) as GR, where artisanal reef fisheries are regulated (via limitations on the use of fishing gear), and the northern GoA regions (South Nuweiba, North Nuweiba and Taba) as OA, where fisheries management appear to be more permissive than other regions and a variety of fishing gears are used (e.g., hook and line, nets, traps, spear guns) throughout these regions (Figure 1; Table 1).

**Figure 1.**
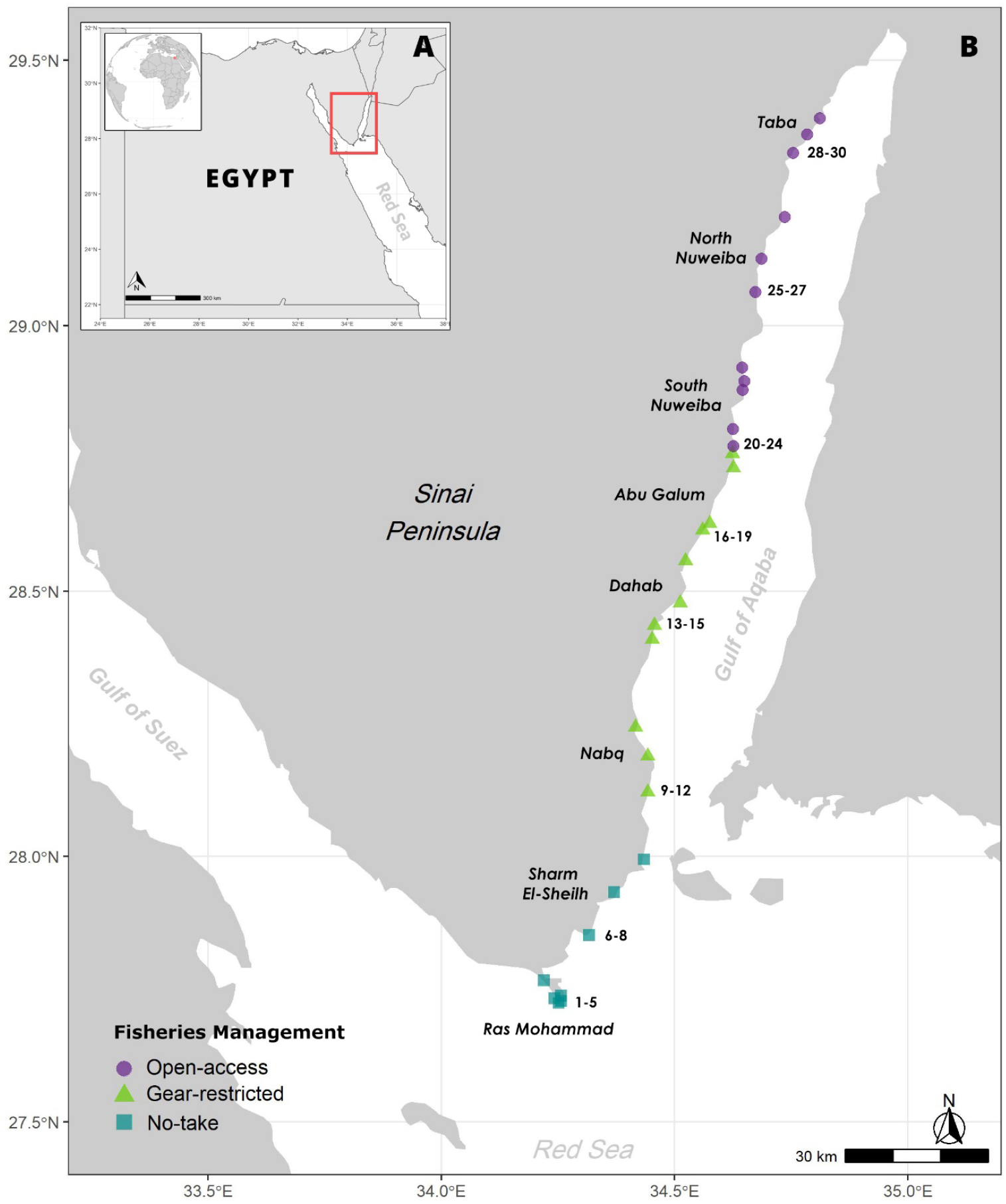
Map of Egypt, showing the location of the GoA (A). Map of the GoA, showing locations of the eight study regions representing 30 reef sites (B). Different shapes represent fisheries management type; (square symbols) represent NT, (triangles) represent GR, and (circles) represent OA reefs. Surveyed sites per region were assigned a numerical code (shown next to regions).

**Table 1.** The surveyed regions were stratified according to fisheries management systems as (no-take, gear-restricted, and open-access). The uses allowed (√), permitted in the designated zones (P), or prohibited (x) within each region are shown. The potential effects of the anthropogenic stressors on coral reefs are also shown. Severity of each impact is shown as: (-) non-significant; (+) noticeable; (++) marked; (+++) severe. Adopted from (Galal, 1999; Ashworth, 2004; Mabrouk, 2015) to include the northern regions of the GoA.

| Local anthropogenic threats | No-take |  | Gear-restricted |  |  | Open-access |  |  |
| --- | --- | --- | --- | --- | --- | --- | --- | --- |
|  | RM | SH | NQ | DB | AG | SN | NN | TB |
| **Fishing**<br>(Gill & Trammel nets, Hand-lining) | x | x | P | P | P | √ | √ | √ |
|  | - | - | ++ | ++ | ++ | +++ | +++ | +++ |
| **Harvesting of reef organisms**<br>(Collection of Shells and Octopus) | x | x | P | P | P | √ | √ | √ |
|  | - | - | +++ | +++ | +++ | +++ | +++ | +++ |
| **Coastal construction**<br>(Camps, Villages, Hotels) | x | √ | P | √ | P | x | √ | √ |
|  | - | +++ | + | +++ | ++ | - | +++ | +++ |
| **Recreational activities**<br>(Snorkeling, SCUBA diving) | √ | √ | √ | √ | √ | x | √ | √ |
|  | +++ | +++ | + | +++ | ++ | - | + | + |

RM= Ras Mohammad, SH= Sharm el-Sheikh, NQ= Nabq, DB= Dahab, AG= Abu Galum, SN= South Nuweiba, NN= North Nuweiba, TB= Taba

### 2.2. Herbivorous fish surveys

Species-level visual censuses targeting four key herbivorous fish families (Acanthuridae, Kyphosidae, Scaridae, and Siganidae) were conducted using standard underwater visual belt-transect survey method. Within each site, four 50 × 10 m belt transects were laid along the reef crest habitat, oriented parallel to the coastline and each was separated by a minimum at least 5 m to ensure independence between transects. In each transect, herbivorous fishes were counted and the body length of each individual was visually estimated to the nearest 5cm. Surveys focused on counting fishes (≥10 cm total length, TL), except for the two small surgeonfish, *Acanthurus nigrofuscus* and *Ctenochaetus striatus* (≥5 cm, TL). Each fish was classified into one of the following nine (5 cm) size classes; 6-10, 11-15, ……, 41-45 and 46-50 cm. Fish density was expressed as number of individuals per 500 m^2^. All surveys were limited to reef crest habitat (1–3 m, depth) and conducted during daylight hours between 10:00 and 17:00 hrs at high tide to ensure that counts were undertaken when the reef crest habitat was available for grazing fishes.

### 2.3. Estimations of fish biomass

Fish densities and length estimates were converted to kilograms (kg) of biomass per unit area (kg/500 m^2^) using the allometric length-weight equation: W= aTL^b^, where W is weight in kilograms, TL is the total length in cm (taking the midpoint of each size class), and parameters a and b are species-specific constants derived for each species from FishBase (Froese and Pauly 2019) and other published sources in the region (Table S2). Herbivore species were placed into one of four functional feeding groups: excavators, scrapers, grazers, and browsers (Green and Bellwood, 2009). However, species that were not categorized in the aforementioned references, such as *Chlorurus sordidus*, was identified as excavator, *Siganus luridus* and *Siganus rivulatus* as macroalgal browsers (Ebrahim *et al*., 2020), *Siganus stellatus* and *Ct. striatus* as grazers (Afeworki et al., 2013; Marshell and Mumby, 2012) (Table S3).

### 2.4. Data analyses

For each fish species, the density, biomass, and frequency of occurrence metrics were calculated as a percentage of the grand total for each metric. Subsequent, an index of relative dominance (IRD) for each species was created by multiplying the percent frequency of occurrence of the species combination by the relative percent biomass of that species (Friedlander *et al*., 2003).

#### 2.4.1 Univariate data analysis

Herbivorous fish characteristics (biomass, density, species richness, body length) among different FMSs were analysed using generalized linear mixed models (GLMMs). All models included the predictors ‘fisheries management’ as (fixed effect with three levels: NT, GR, OA), while the two spatial variables (region, site) were as random effects to account for their potential influence upon herbivorous fish characteristics, as response variables. Mean herbivorous fish biomass and body length were modelled using a Gamma distribution (R package lme4) (Bates *et al*., 2015), which is appropriate for continuous data (Zuur *et al*., 2009), while species richness and numerical density (abundance) were modelled using a Poisson distribution (R package lme4), which is appropriate for count data (Zuur *et al*., 2009). All models were fitted with a “log” link function. A Tukey post-hoc test was used to investigate pairwise differences between significant factor levels using the *glht* function of the multcomp package (Hothorn *et al*., 2008) in R project 4.0.3 (R Core Team 2020).

#### 2.4.2 Multivariate data analysis

(A). To examine variability in herbivorous fish biomass and density between FMSs, we used multivariate statistical and ordination methods implemented through the R package vegan (Oksanen et al., 2019). At site level, fish species density and biomass for the entire herbivorous fishes and for each individual feeding group were averaged and distance matrices based on Bray–Curtis dissimilarities were created using the *vegdist* function. First, we conducted permutational multivariate analysis of variance (PERMANOVA) tests using the *adonis* function to test whether the biomass and density varied significantly among the different types of FMSs. Similarly, a permutational analysis of multivariate dispersion (PERMDISP; *betadisper* function) within the FMSs was conducted. Adonis Pairwise comparisons were tested using the *pairwise*.*adonis* function (Martinez 2020). Next, we performed a multivariate ordination (principal coordinates analysis; PCoA) using the *betadisper* function to display the variation observed in PERMANOVA tests. Statistical significance of the PCoA ordinations was determined by an ANOVA-like permutation test (*permutest* function). Finally, to determine the species responsible for driving any observed differences in the herbivorous fish density and biomass, we used the *simper* (similarity percentages) function to identify influential species among FMSs. A cut-off criterion was applied to allow identification of a subset of species whose cumulative percentage contribution reached 70% of the dissimilarity value. The aforementioned analyses were conducted using R version 4.0.3 (R Core Team 2020).

(B). To evaluate whether the mean density and biomass of total herbivorous fish were affected by fisheries management, region or site factors, we used a PERMANOVA (Anderson, 2014), based on a Bray–Curtis resemblance matrix with square-root transformed data. The design was based on three factors: (1) FMSs (fixed factor), (2) region (nested in FMSs, random factor), and (3) site (nested in region and FMSs, random factor). The relative importance of each factor was quantified through the estimates of component of variation. Next, the potential effect of multivariate dispersion was assessed using PERMDISP (Anderson, 2006). Multivariate patterns were visualized through (canonical analysis of principal coordinates; CAP) (Anderson and Willis, 2003) with herbivorous fish density and biomass data of all replicates. In each CAP biplot, correlation vectors based on Spearman ranking (>0.5) were overlaid on the CAP plots to determine which fish species (vectors) were responsible for the clusters. Analyses and CAP ordination were performed in using PRIMER 6.1.13 and in the PERMANOVA + 1.0.3 add-on (Clarke & Gorley 2001, Anderson et al. 2008).

## 3. Results

### 3.1. Herbivore assemblages

Overall, 120 belt-transects covering an area of 60 km^2^ were performed on reef crest zone across 30 study sites. A total of 12,542 fish belong to 24 herbivorous species (8 browsing, 3 excavating, 6 grazing, and 7 scarping) were recorded (Table S3). One-third of the observed herbivore species (10 spp.) were widely distributed, occurring on 80-100% of survey sites (Table S4). Among the encountered species, only two were endemic to the Red Sea region (Golani and Fricke, 2018): *C. gibbus* and *C. viridescens*, together comprised only 2.2 and 3.5% of the total density and total biomass, respectively (Table S4). The most dominant species based on IRD was *A. sohal*, which was present in 55 % of transects and accounting for one-quarter of the total biomass. The next most dominant species were *C. sordidus*, and *S. niger*, which together accounted for 18.1% of the total biomass and 14.6% of the total density (Table S4). The overall abundant species in terms of numerical density were the small-bodied *A. nigrofuscus* and *C. striatus* grazers, together accounting for 45.4% of the total density. The most dominant functional group was grazers (61.2 and 37.8% of total density and biomass, respectively), followed by browsers (16.3% and 22.6%), scrapers (12.6% and 26.5%), and excavators (9.9% and 13.0%).

### 3.2. Total herbivorous fish

The grand mean biomass of herbivores at unfished NT reefs with 36.2±2.1 kg (per 500 m^2^) and moderately fished GR reefs with 23.7±4.0 kg was significantly greater by ∼4.3 and 2.8 times than in heavily fished OA reefs (GLMMs, *t* = -6.65, *P*<0.001; pairwise test, *P*<0.001; Figure 2a; Tables 2, 3). Biomass values varied considerably among regions and across sites, ranging from 2.9±0.5 kg at heavily fished OA sites of Taba, to 38.4±1.8 kg in unfished NT sites of Ras Mohammed (Table S5a). Among GoA regions, the unfished regions (Ras Mohammed and Sharm El-Sheikh) and only one of fished regions (Nabq) met the global mean herbivore biomass from unfished reefs of ∼30 kg/500 m^2^ (Edwards *et al*., 2014). (Figure S1; Table S5a). Comparing reef site-level biomass to a global mean herbivore biomass target, we found that 87.5% of NT sites exceeded this target, compared to 27.3% of GR sites. Noticeably, none of the OA sites have reached this target (Table S5a).

**Figure 2.**
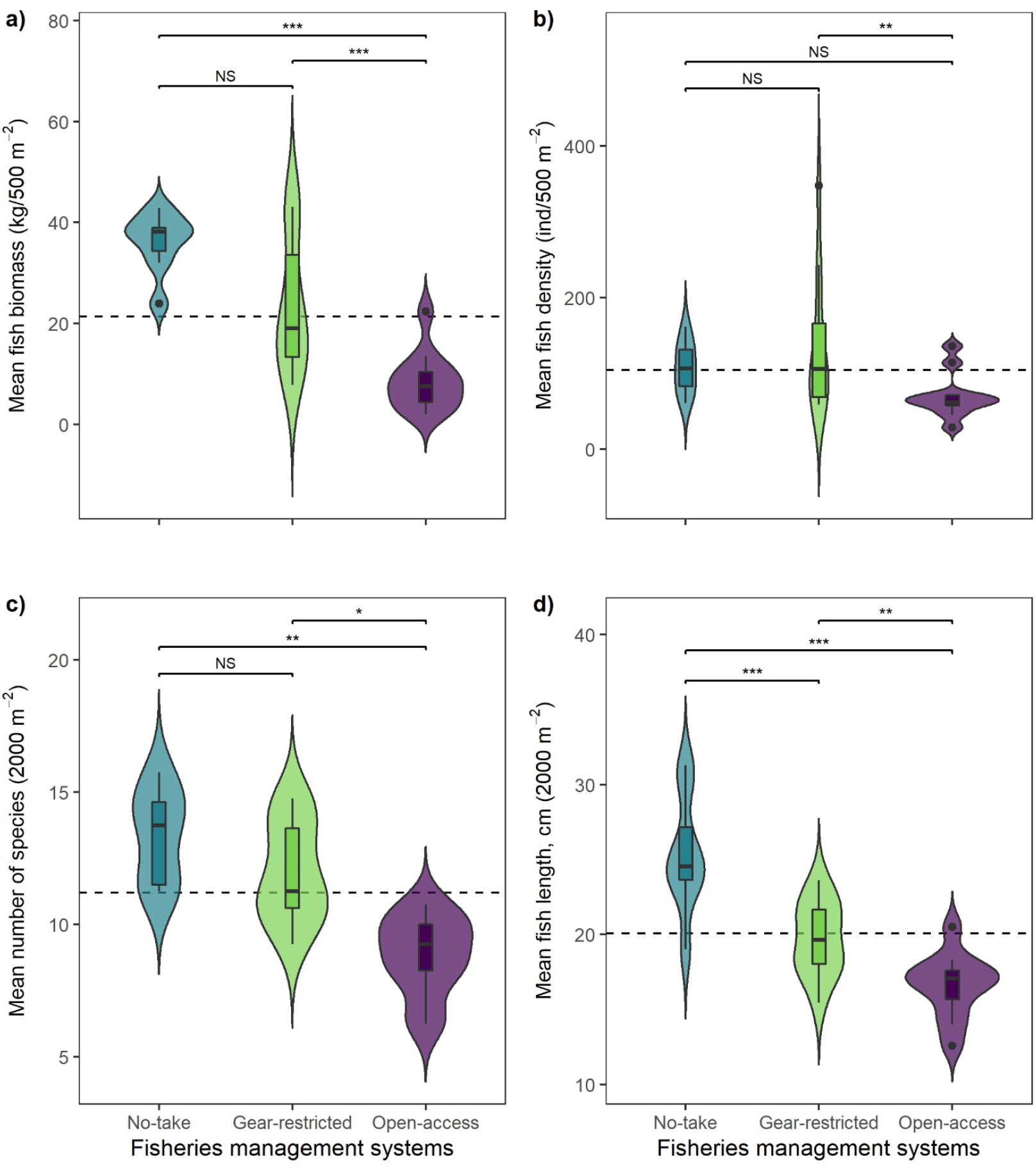
Violin plots with boxplots overlaid, showing the comparison of total herbivorous fish indices between the sampling three fisheries management systems in 2017. Fish biomass (a), Fish density (b), Species richness (c) and Fish size (d). The boxplot shows the median (black line) and interquartile range (top and bottom borders of the box). Tukey post-hoc test used to examine differences in management levels: levels of significance is denoted with asterisks \**P*<0.05, \*\**P*<0.01, \*\*\**P*<0.001, NS non-significant. Dashed horizontal line represent the overall mean value for each fish index combined across all management systems.

**Table 2.** Summary of generalized linear mixed-effects models (GLMMs) results used to examine the differences in fish biomass, density, species richness and body length. Models contain the fixed effect: fisheries management systems (no-take, gear-restricted, and open-access). SE = standard error, Gamma distribution models use a *t* value while Poisson models use a *z* value.

| Response variable | Predictor variable | Estimate | SE | *t/z* value | *p* value |
| --- | --- | --- | --- | --- | --- |
| Biomass | Intercept | 4.9610 | 0.1884 | 26.333 | < 0.001 *** |
|  | Open-access | -1.6455 | 0.2476 | -6.646 | < 0.001 *** |
|  | Gear-restricted | -0.5543 | 0.2476 | -2.239 | 0.025 * |
|  | Intercept | 2.5311 | 0.4740 | 5.339 | < 0.001 *** |
|  | Open-access | -0.6818 | 0.6129 | -1.112 | 0.266 |
|  | Gear-restricted | -0.1055 | 0.6125 | -0.172 | 0.863 |
|  | Intercept | 3.5700 | 0.2540 | 14.053 | < 0.001 *** |
|  | Open-access | -1.6126 | 0.3339 | -4.830 | < 0.001 *** |
|  | Gear-restricted | -0.5357 | 0.3339 | -1.605 | 0.109 |
|  | Intercept | 3.9424 | 0.2972 | 13.267 | < 0.001 *** |
|  | Open-access | -1.9482 | 0.3905 | -4.988 | < 0.001 *** |
|  | Gear-restricted | -0.8473 | 0.3905 | -2.170 | 0.030 * |
|  | Intercept | 3.1213 | 0.2792 | 11.178 | < 0.001 *** |
|  | Open-access | -1.6041 | 0.3668 | -4.374 | < 0.001 *** |
|  | Gear-restricted | -0.2279 | 0.3666 | -0.622 | 0.534 |
| **Density** |  |  |  |  |  |
|  | Intercept | 6.0105 | 0.1605 | 37.452 | < 0.001 *** |
|  | Open-access | -0.4519 | 0.2125 | -2.127 | 0.033 * |
|  | Gear-restricted | 0.1220 | 0.2106 | 0.579 | 0.562 |
|  | Intercept | 3.6468 | 0.1770 | 20.606 | < 0.001 *** |
|  | Open-access | -0.2538 | 0.2366 | -1.072 | 0.284 |
|  | Gear-restricted | 0.1138 | 0.2311 | 0.492 | 0.622 |
|  | Intercept | 4.2388 | 0.3047 | 13.910 | < 0.001 *** |
|  | Open-access | -1.2588 | 0.4021 | -3.131 | 0.002 ** |
|  | Gear-restricted | -0.2462 | 0.3948 | -0.624 | 0.533 |
|  | Intercept | 5.4232 | 0.1944 | 27.900 | < 0.001 *** |
|  | Open-access | -0.3017 | 0.2557 | -1.180 | 0.238 |
|  | Gear-restricted | 0.1192 | 0.2555 | 0.467 | 0.641 |
|  | Intercept | 3.8109 | 0.2819 | 13.517 | < 0.001 *** |
|  | Open-access | -0.3978 | 0.3664 | -1.086 | 0.278 |
|  | Gear-restricted | 0.5423 | 0.3635 | 1.492 | 0.136 |
| **Species richness** |  |  |  |  |  |
|  | Intercept | 2.93784 | 0.08138 | 36.101 | < 0.001 *** |
|  | Open-access | -0.35213 | 0.11607 | -3.034 | 0.002 ** |
|  | Gear-restricted | -0.06273 | 0.10840 | -0.579 | 0.563 |
| **Fish size** |  |  |  |  |  |
|  | Intercept | 3.22531 | 0.04587 | 70.319 | < 0.001 *** |
|  | Open-access | -0.42615 | 0.06028 | -7.070 | < 0.001 *** |
|  | Gear-restricted | -0.25130 | 0.06028 | -4.169 | < 0.001 *** |

Values in boldface type are statistically significant at $\alpha = 0.05$ , and asterisks indicate levels of significance \* $P < 0.05$ , \*\* $P < 0.01$ , \*\*\* $P < 0.001$

**Table 3.** Results of multiple comparisons of means (Tukey Contrasts) between levels of fisheries management systems for each response variables.

| Response variable | Fisheries Management Systems |  |  |
| --- | --- | --- | --- |
|  | NT vs OA | GR vs OA | NT vs GR |
| **Biomass** |  |  |  |
| All herbivores | < 0.001 *** | < 0.001 *** | 0.065 |
| Excavators | 0.506 | 0.545 | 0.984 |
| Scrapers | < 0.001 *** | 0.001 ** | 0.243 |
| Grazers | < 0.001 *** | 0.006 ** | 0.076 |
| Browsers | < 0.001 *** | < 0.001 *** | 0.808 |
| **Density** |  |  |  |
| All herbivores | 0.084 | 0.009 ** | 0.831 |
| Excavators | 0.531 | 0.213 | 0.875 |
| Scrapers | 0.005 ** | 0.015 * | 0.807 |
| Grazers | 0.465 | 0.172 | 0.887 |
| Browsers | 0.522 | 0.013 * | 0.294 |
| **Species richness** |  |  |  |
| All herbivores | 0.007 ** | 0.022 * | 0.831 |
| **Fish size** |  |  |  |
| All herbivores | <0.001 *** | 0.005 ** | < 0.001 *** |

NT refers to No-take, GR refers to Gear-restricted and OA refers to Open-access. Values in boldface type are statistically significant at $\alpha = 0.05$ , and asterisks indicate levels of significance \* $P < 0.05$ , \*\* $P < 0.01$ , \*\*\* $P < 0.001$

The grand mean density of herbivores in GR reefs with 136.5±27.6 individuals (per 500 m^2^) was higher by 1.3 and 1.9 times than at NT reefs and OA reefs, respectively, though not significant (GLMMs; *P*>0.05; Figure 2b; Tables 2, 3). Density varied considerably among regions and across sites, ranging from 48.2±10.0 ind. at heavily fished OA sites of Taba, to 188.1±59.4 ind. at moderately GR sites of Nabq (Figure S1; Table S5b). Species richness of herbivores declined significantly with increasing fishing pressure, from 16.9±1.1 species (per 2000 m^2^) at NT reefs to 11.3±0.5 species at OA areas (GLMMs; *P*=0.002; Figure 2c, Table 2). Notably, six species (four browsers, one grazer and one excavator) were not observed at OA sites. A pairwise comparison between NT reefs and GR reefs revealed non-significant variation (pairwise test, *P*=0.83, Table 3). At region level, the highest species richness was recorded at Ras Mohammed 19±0.7 species and the lowest at Taba 9.7±0.3 species (Figure S1; Table S6a).

In accordance with the biomass variation, herbivorous fish size declined significantly from 25.4±3.9 cm (per 2000 m^2^) at NT reefs to 19.7±2.5 cm in GR reefs, and to 16.6±2.2 cm at OA reefs (*P*<0.001 for all comparisons; Figure 2d, Table S6b). Size-class distribution of herbivorous fishes in fished (GR and OA) reefs were dominated by small-sized individuals (6–20 cm in length) with 77.4% and 81.1% of the total density, representing 30.8% and 37.3% of the total biomass, respectively. However, a moderate frequency of small-sized individuals was recorded in unfished NT reefs with 50.8% of the total density representing 12.2% of the total biomass (Figure S2). In marked contrast, only 8.8% and 3.5% of the total density on fished (GR and OA) reefs were larger than 30 cm, compared to 26.1% at NT reefs. The variation is even greater in terms of biomass, with NT reefs having more than 45 times the biomass of large (> 35 cm) herbivores when compared with heavily fished OA reefs (Figure S2).

### 3.3. Functional groups

Biomass, species richness and size-frequency distributions of the four herbivore functional groups declined considerably with increasing fishing pressure on the GoA (Figure 3; Tables S5 and S6). These three herbivore metrics were all generally lower at heavily fished OA reefs and moderately fished GR reefs compared to unfished NT reefs (Figures 3, S4; Tables S5, S6). Three functional groups (scrapers, grazers, browsers) revealed significant declines in biomass with increasing fishing pressure. This pattern was most notable when comparing NT with OA reefs (GLMMs; *P*<0.001 for the three functional groups; Figure 3; Tables 2 and 3). However, the biomass of excavators showed a little variation among the investigated FMSs except for NT reefs where they showed considerably higher abundance for size-classes >35 cm (Figure 3). Size-class distribution of excavators declined from 27.5±1.2 cm (per 2000 m^2^) at NT reefs to 24.5±0.5 cm in GR reefs, and to 23.4±0.6 cm at OA reefs (Table S6b). Furthermore, excavators at fished (GR and OA) reefs were generally dominated by small-sized individuals (6–20 cm in length) with 77.4% and 81.1% of the total density compared to 50.8% at unfished NT reefs (Figure S3). Notably, large-sized excavators (>40 cm) were totally absent from OA reefs, distinguishing this assemblage from that on NT reefs (Figures 3, S3).

**Figure 3.**
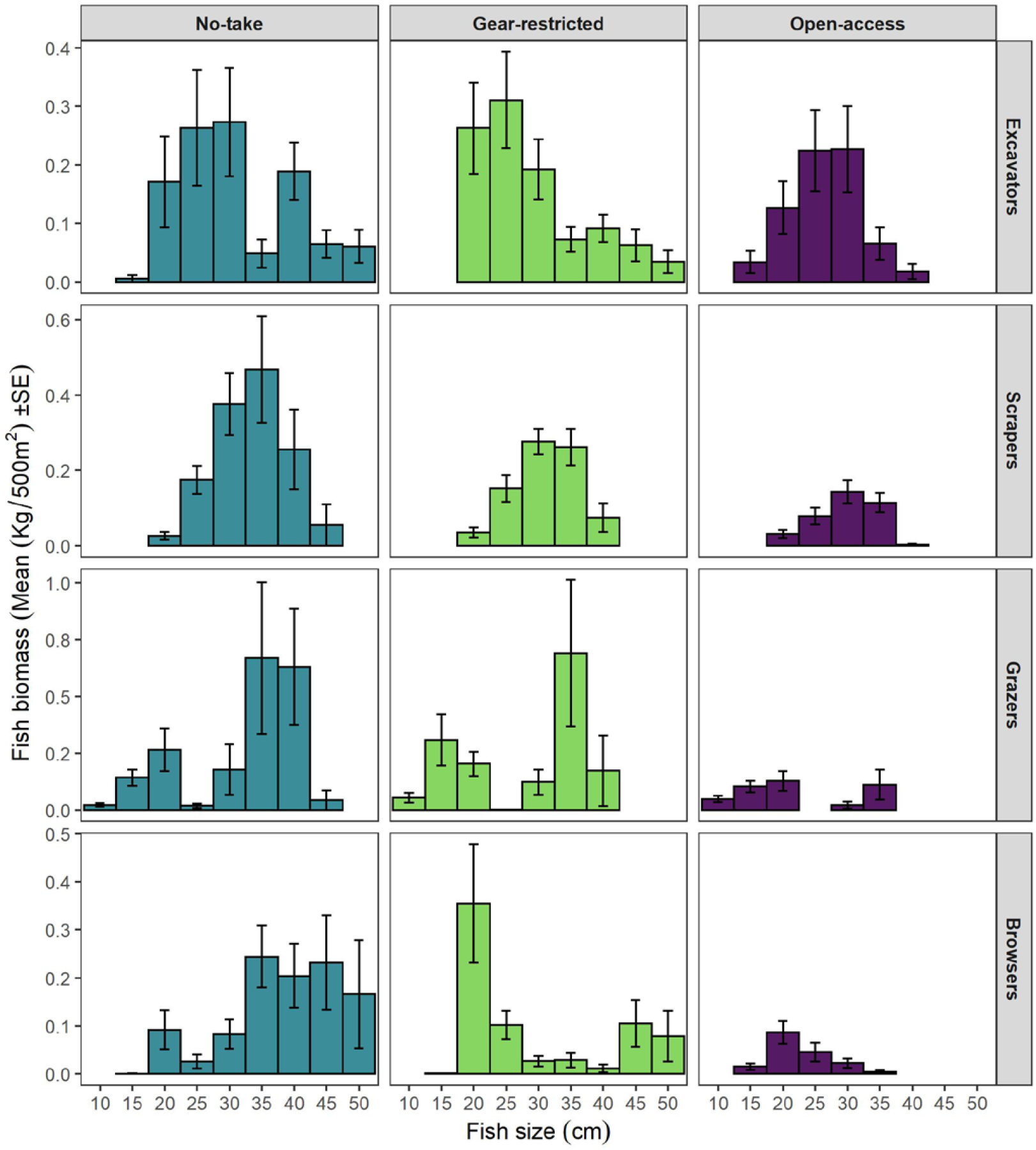
Mean biomass (kg 500^2^ ± SE) for each size-class (total length; cm) of herbivorous fishes divided by (functional feeding group) in each fisheries management systems

In respect to scrapers, biomass and density varied significantly between NT and OA reefs (GLMMs: *P*<0.001 for all comparisons; Figure 3, S4; Table 2). Size-class distribution of scrapers declined considerably from 31.9±1.0 cm at NT reefs to 27.8±0.4 cm in OA reefs (Table S6b). Among grazers and browsers, the biomass results revealed high significant differences between NT reefs and OA reefs. On the other hand, density revealed no significant differences (GLMMs: *P*<0.001 for all comparisons; Figure 3, S4; Tables 2, 3). However, differences in mean body-length of grazers and browsers between individual FMSs revealed that grazers and browsers were found to be considerably larger on NT reefs than on OA reefs (Figure 3, S4; Table S6b). Both functional groups were dominated by medium and small sized fishes on all reefs. At NT reefs, most of the grazers and browsers biomass came from the large-sized classes (more than 30 cm), contrary to GR and OA reefs were in the size classes less than 30 cm (Figure 3).

At region level, Mean biomass and density of excavators were highest at Abu Galum with 3.7±0.4 kg (per 500 m^2^) and 13.3±2.0 individuals (per 500 m^2^), respectively, followed by Ras Mohammed and Sharm El-Sheikh (Figure S4a, Table S5a). Biomass and density values of scrapers varied greatly among regions, where Ras Mohammed having the highest value 12.2±2.6 kg and 23.6±6.3 ind., whereas Taba had the lowest biomass value 0.6±0.2 kg and North Nuweiba had the lowest density with 2.9±1.4 ind. Notably, grazers and browsers biomass and density were considerably higher in Nabq with (18.1±4.7 and 9.5±2.5 kg) and (107.2±33.5 and 52.2±24.2 ind.) respectively, and the lowest values were at heavily fished OA regions (Figure S4b, Table S5b).

### 3.4. Herbivorous fish community structure

#### 3.4.1. Effects of fisheries management on fish biomass and density

The total biomass of herbivorous fish communities varied significantly between FMSs (*R*^*2*^=0.32, *P*=0.001; Table S7). SIMPER results showed that over 70% of the dissimilarity in herbivore community structure in NT compared to the OA fishery was attributable to the highest biomass of 7 species. In particular, the biomass of the three large-bodied herbivores (the grazer *A. Sohal*, the algal browser *N. unicornis*, and the scraper *H. harid*) contributed most to dissimilarities (53.5%) (Table S9). The biomass structure of total herbivores, and each individual functional group was also significantly influenced by FMSs (Figures 4a-d, S5; Table S5). In OA reefs, the decline in herbivorous biomass (either total or functional groups) was mostly driven by the declines in body length (Figures 3, S3; Table S5), particularly the grazer (*A. sohal*), scraper (*H. harid*) and macroalgal browsers (*N. unicornis, N. elegans*) (Table S9).

**Figure 4.**
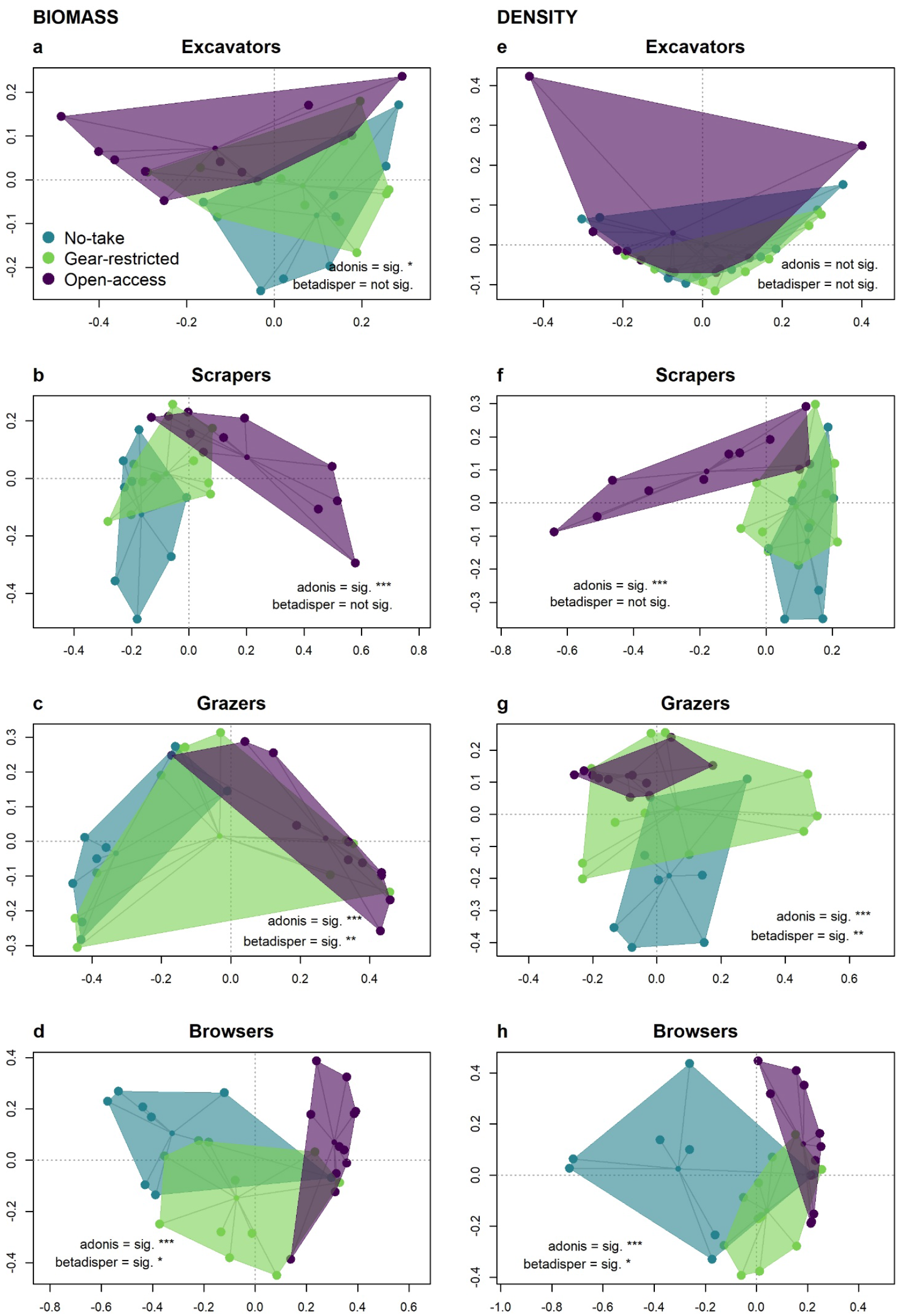
Principal coordinates analysis (PCoA) ordination of distances among the centroids of main effects (fisheries management) based on Bray–Curtis dissimilarities of biomass (left column) and density (right column) for individual herbivores groups; excavators (a, e), scrapers (b, f), grazers (c, g), and browsers (d, h). Points represent individual sites (connected to the centroid point in the center), sites are color-coded by fisheries management systems, and shaded polygons indicate boundaries of observed community structure. Significance level of ADONIS and PERMDISP tests are denoted by \*\*\**P*<0.001, \*\**P*<0.01, \**P*<0.05.

The total density of herbivorous fish communities also varied significantly across FMS (*R*^*2*^=0.25, *P*=0.001; Table S7). In terms of functional groups, all groups except for the excavators exhibited significant differences in structure across FMSs (Figures 4e–h, S5; Table S5). SIMPER results showed that over 70% of the dissimilarity in herbivore community structure in the NT compared to the OA fishery was attributable to the highest numerical density of 6 species (53.1%) in NT reefs and 2 species (20.7%) in OA reefs (Table S10). The density of the four large-bodied herbivores (the grazer *A. Sohal*, the scrapers *H. harid* and *S. ferrugineus*, and the excavator *C. sordidus*) contributed most to the dissimilarities (32.6%) between NT and OA reefs. Conversely, the influential small-bodied grazer species *A. nigrofuscus* and the algal browser *S. luridus* experienced increases in density in GR and OA reefs (Table S10).

#### 3.4.2. Effects of site, region and fisheries management on fish biomass and density

CAP analysis showed that FMSs effects are the strongest, especially contrasting NT versus either GR or OA along CAP axis-1, in addition to the significant differences between FMSs in terms of fish biomass and density (*P*=0.0001, Figure 5a, b). Unfished NT reefs are characterized by greater biomass of highly targeted fish browsers (*K. cinerascens, N. elegans, N. unicorn*), grazers (*A. sohal*) and scrapers (*H. harid, S. ferrugineus*), recording 5.8-32.9 and 1.9-5.7 times greater than the OA and GR reefs, respectively. However, the heavily fished OA reefs have a greater biomass and density of the only scarid browsers (*C. viridescens*). Lastly, moderately fished GR reefs were distinct from other management types by having a greater biomass and density of siganid browsers (*S. luridus* and *S. rivulatus*). The observed patterns in CAP ordination were confirmed by results from PERMANOVA test, which showed that, fish biomass and density significantly differed among FMSs, regions, and sites (*P<0*.*05* for all comparisons; Table S8). For biomass and density, the estimates of component variation (ECV) test in PERMANOVA indicated that site variation was greater followed by fisheries management effects (Table S8).

**Figure 5.**
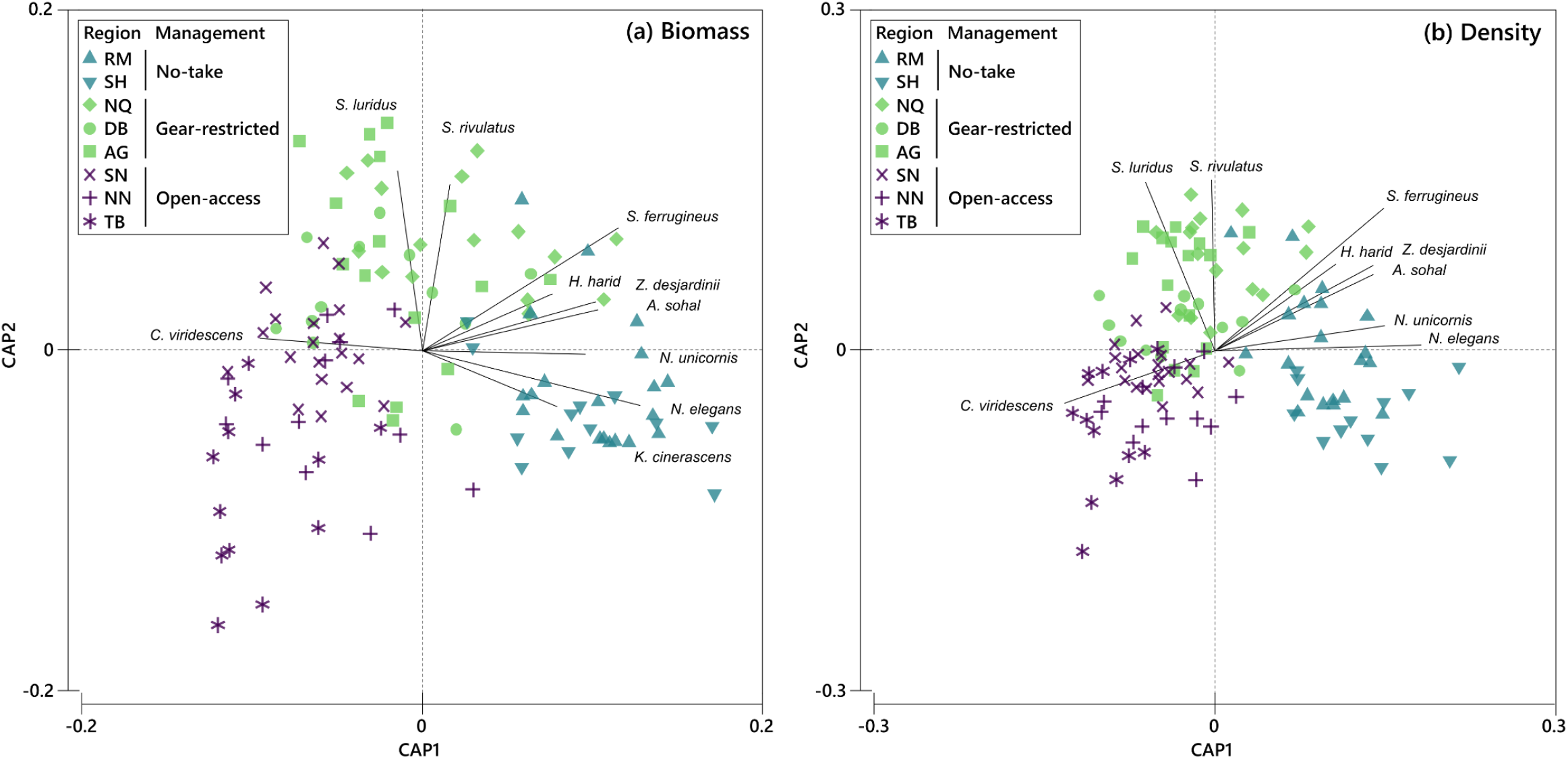
Canonical analysis of principal coordinates (CAP) ordination based on Bray-Curtis similarity for herbivorous fish biomass (a) and density (b) across three fisheries management systems (NT, GR, OA) and eight regions (RM: Ras Muhammad, SH: Sharm El-Sheikh, NQ: Nabq, DB: Dahab, AG: Abu Galum, SN: South Nuweiba, NN: North Nuweiba, TB: Taba). Each color represents a management type and each shape a study region. Species correlations with the canonical axis are represented as vectors for species with Pearson R value greater than 0.5; e.g. biomass and density of large-bodied algal browsers (*N. unicornis* and *N. elegans*) are positively correlated with NT management. Choice of m = 8 for fish biomass data (Eigenvalues of correlation: 1 = 0.85, 2 = 0.65; Permutation test: *P* = 0.0001) and m = 10 for fish density data (Eigenvalues of correlation: 1 = 0.86, 2 = 0.69; Permutation test: *P* = 0.0001).

## 4. Discussion

MPAs managers challenge the issues of balancing the ever-increasing demand for coral reef fisheries with maintaining reef health and resilience. The effective management of herbivorous fish populations is one tool that reef managers can use to complement existing management strategies to improve coral reef resilience. In South Sinai (northern Red Sea), the existing MPAs are subject to different levels of fishing pressure, therefore, understanding the effects and consequences of this stressor on herbivorous fish populations is highly needed to properly manage current and future levels of fishing.

In this study, we found that herbivorous fish assemblages at locations closed to fishing (NT) support on average more than 1.5 and 4 folds the total biomass, compared to those with restrictions on fishing gears (GR), and open to all fishing gears (OA), respectively. These results are consistent with the growing number of regional and global studies that noted similar patterns between unfished and fished areas (Edwards *et al*., 2014; Advani *et al*., 2015; Kattan *et al*., 2017; Steneck *et al*., 2018; Campbell *et al*., 2018, 2020; Rolim *et al*., 2018; Bejarano *et al*., 2019; Cinner *et al*., 2020; Humphries *et al*., 2020; Foo *et al*., 2021). For example, Edwards et al (2014) examined the impacts of fishing on herbivorous fishes from 145 locations around the world. They found that biomass is more than twice as high in locations closed to fishing relative to fished locations. Campbell et al (2018) observed the same patterns within 22 global marine ecoregions. They showed that herbivorous fish (Scaridae and Siganidae) biomass in unfished reefs were (46% and 330% higher, respectively) than heavily fished reefs. Furthermore, recent findings by Campbell et al (2020) corroborate our results that NT and GR reefs were associated with significantly higher biomass of herbivores relative to OA reefs. Cinner et al (2020) found that gear-restricted MPAs provided similar conservation benefits as fully protected MPAs for parrotfish family, and thus, for this particular group of fish, gear restrictions can be very effective. To best of our knowledge this is the first study to investigate the variation patterns of biomass and population size structure of herbivorous reef fishes between fished and unfished locations in the GoA (northern Red Sea).

Similar to the observed patterns in total herbivorous fish biomass and body length, species richness also declined with increasing fishing pressure and was considerably higher in unfished reefs (NT). This finding is consistent with previous studies that reported a positive effect of protection on herbivorous fish species richness (Bejarano *et al*., 2019; Topor *et al*., 2019; Altman-Kurosaki *et al*., 2021). Despite consistent patterns in herbivorous fish sizes, biomass and richness, there was no significant difference in numerical density between fished (GR and OA) and unfished (NT) locations. Previous studies which investigated fishing impacts on the abundance of herbivorous fish populations offer conflicted opinions. Several authors have concluded that fishing reduced herbivorous fish abundances e.g. (McClanahan *et al*., 1999; Edwards *et al*., 2014; Russ *et al*., 2015; Steneck *et al*., 2018). Whereas, observations made by other studies suggested that herbivorous fish abundances, in particular parrotfish family appeared to be more influenced by protection status (Price *et al*., 2021), environmental conditions (McClanahan and Muthiga, 2020), and habitat characteristics (Vallès and Oxenford, 2014) rather than fishing effect. On the other hand, density of surgeonfish species were influenced more by the changes of benthic composition rather than protection status (Russ *et al*., 2018). However, small sized individuals and the small-bodied fish (surgeonfish, rabbitfish and parrotfish) species increased with increasing fishing pressure (Hawkins and Roberts, 2004; Clua and Legendre, 2008; Bellwood *et al*., 2012; Advani *et al*., 2015; Bejarano *et al*., 2019). Many of the latter observations would appear to corroborate the predator-removal hypotheses developed by Boaden and Kingsford (2015). In our study, the numerical density of herbivorous fish assemblages appeared to be driven by the highly abundant surgeonfish family (Acanthuridae), particularly by the small-bodied grazer *A. nigrofuscus*, which is considerably more abundant in fished locations (GR and OA). Similarly, abundances of the highly targeted rabbitfish family (Siganidae), particularly *S. luridus* and *S. rivulatus*, were also more abundant in fished locations. Our findings support those of Floeter et al (2006), who also found highly targeted herbivore species were significantly more abundant and larger in a highly protected sites.

Predatory species have been recognized as important in maintaining diversity on coral reefs (Hixon, 2015). In the GoA, groupers family (Serranidae) recognised to heavily prey on some herbivores, particularly small acanthurids (Shpigel and Fishelson, 1989), and so, it is likely that interactions may explain the higher density of small-bodied grazers (surgeonfish) in fished areas compared to unfished areas (Ashworth and Ormond, 2005). Neither species of rabbitfish family (Siganidae) are considered to be prey species of groupers (Shpigel and Fishelson, 1991), Accordingly, the predator-removal hypothesis mayn’t considered as a factor that determine the higher abundances observed in fished areas (e.g. Nabq). However, this study suggests that the increasing of abundance of rabbitfish family might be affected more by environmental factors as mentioned by (Olds *et al*., 2013; Roff *et al*., 2019) rather than predation or fishing effect. Price et al (2021) found that abundances of grazers and browsers increased in the fished areas with high algal cover. Personal observations suggest the presence of seagrass beds and/or mangroves close to fished reefs (particularly at Nabq), could be an important factor in determining the significance of Siganids associated with them.

Most of fisheries tend to target commercially important reef fishes with larger sizes (Wilson *et al*., 2010; Robinson *et al*., 2017). Handline, gillnets and trammel nets are the more widely used traditional fishing methods by Bedouin fishermen along the Egyptian GoA. However, in the last decade, spearfishing has become more frequently practiced by Bedouin (Poonian, 2020). This highly selective method contributes to removing larger size classes and key herbivorous fish species (Frisch *et al*., 2012; Bender *et al*., 2014; Barbosa *et al*., 2021). In many regions around the world, the selectivity of larger herbivore species has been observed (Edwards *et al*., 2014; Ford *et al*., 2016), but this is the first study to demonstrate this pattern in the GoA for this fish group. The striking depletion of large-bodied herbivorous fish density and biomass (*A. sohal, H. harid, N. elegans, S. ferrugineus*; > 35 cm TL), was noticeable by the drop from 12.9 and 36.4% of the total density and biomass, respectively on NT reefs to 0.1 and 0.8%, respectively on OA reefs. Moreover, large-bodied target species (*C. bicolor, N. unicorn, Kyphosus* spp.) were virtually absent from OA reefs.

As large-bodied herbivore species have important and unique ecological function in maintaining the health and resilience of coral reefs, hence, protecting and restoring these key herbivores has become a priority (Heenan and Williams, 2013; Mumby *et al*., 2016). Furthermore, conserving the functional redundancy and complementary roles of diverse herbivorous fish community are critical for maintaining healthy coral reef ecosystems (Bellwood *et al*., 2006, 2012; Burkepile and Hay, 2008; Cheal *et al*., 2010). Our findings revealed that many of these key herbivore species in the have been overexploited central-northern regions of the GoA (e.g. Dahab, Nuweiba and Taba) and thus may explain the significant increases in turf algae and macroalgae cover that have been reported recently in Dahab (Naumann *et al*., 2015; Reverter *et al*., 2020). We believe that more attention should be paid for managing and maintaining the biomass and composition of these key herbivorous fish groups to improve the existing fisheries management strategies in SSMPAs. Collectively, our findings suggest that if the current fishing level in the central-northern GoA reefs are not effectively managed, it could reduce herbivore communities’ ability to maintain top-down control on algal communities and could lead to degradation of coral communities.

### 4.1. Conclusion

The status of herbivorous reef fishes in the northern Red Sea and Gulf of Aqaba has not been evaluated on such a large scale, so our results provide important baselines for herbivorous fish assemblages in this region. The biomass of herbivorous fishes was 4.3 times higher on unfished reefs than in locations completely open for fishing at the northern Gulf of Aqaba (Nuweiba and Taba). While fishing has a clear effect on biomass, species richness and body length, there is no difference in the numerical density of herbivores between fished and unfished reefs. This contrary pattern proves that larger-bodied individuals and species were excessively impacted by fishing. As a result, our findings highlighting that current levels of fishing (i.e., selectivity of large-bodied herbivores) in the northern Gulf of Aqaba may reduce the ability of grazing fishes to control turf and macroalgae that are growing on the reef substratum and, indirectly, reduce the suitable substrate for coral settlement. Finally, our findings support previous studies showing that fishing-gear restriction provides comparable protection benefits to herbivore groups as no-take MPAs. Therefore, we suggest that such an approach would help coral reef managers to give the overexploited herbivorous fisheries in Nuweiba and Taba a chance to recover.

## Acknowledgments

We thank the South Sinai MPAs sector for administrative support and cooperation to carry out the research. We would like to thank Ismael Abdel-Azim and Ahmed El-Sadek for their help in field work, and Eslam Osman for his valuable help in data analysis. This work is part of the PhD thesis of the first author.

## Data Availability Statement

The data that support the findings of this study are available from the corresponding author upon reasonable request.

## Supporting Information

Additional supporting information may be found online in the Supporting Information section.

## References

Advani S, Rix LN, Aherne DM, Alwany MA, Bailey DM. 2015. Distance from a fishing community explains fish abundance in a no-take zone with weak compliance. PLoS ONE 10: 1–17.

Afeworki Y, Videler JJ, Bruggemann JH. 2013. Seasonally changing habitat use patterns among roving herbivorous fishes in the southern Red Sea: The role of temperature and algal community structure. Coral Reefs 32: 475–485.

Altman-Kurosaki NT, Smith CM, Franklin EC. 2021. O’ahu’s marine protected areas have limited success in protecting coral reef herbivores. Coral Reefs 40: 305–322.

Alwany MA, Thaler E, Stachowitsch M. 2009. Parrotfish bioerosion on Egyptian Red Sea reefs. Journal of Experimental Marine Biology and Ecology 371: 170–176.

Anderson MJ. 2006. Distance-Based Tests for Homogeneity of Multivariate Dispersions. Biometrics 62: 245–253.

Anderson MJ. 2014. Permutational Multivariate Analysis of Variance (PERMANOVA). Wiley StatsRef: Statistics Reference Online: 1–15.

Anderson MJ, Willis TJ. 2003. CANONICAL ANALYSIS OF PRINCIPAL COORDINATES: A USEFUL METHOD OF CONSTRAINED ORDINATION FOR ECOLOGY. Ecology 84: 511– 525.

Ashworth JS. 2004. The effects of protected area status on fish and mollusc stocks in South Sinai, Egypt. Philosophy: 348.

Ashworth JS, Ormond RFG. 2005. Effects of fishing pressure and trophic group on abundance and spillover across boundaries of a no-take zone. Biological Conservation 121: 333–344.

Barbosa MC, Luiz OJ, Cordeiro CAMM, Giglio VJ, Ferreira CEL. 2021. Fish and spearfisher traits contributing to catch composition. Fisheries Research 241.

Bates D, Mächler M, Bolker BM, Walker SC. 2015. Fitting linear mixed-effects models using lme4. Journal of Statistical Software 67.

Bejarano S, Pardede S, Campbell SJ, Hoey AS, Ferse SCA. 2019. Herbivorous fish rise as a destructive fishing practice falls in an Indonesian marine national park. Ecological Applications 29.

Bellwood DR, Hughes TP, Hoey AS. 2006. Sleeping Functional Group Drives Coral-Reef Recovery. Current Biology 16: 2434–2439.

Bellwood DR, Hoey AS, Hughes TP. 2012. Human activity selectively impacts the ecosystem roles of parrotfishes on coral reefs. Proceedings of the Royal Society B: Biological Sciences 279: 1621– 1629.

Bender MG, Machado GR, De Azevedo Silva PJ, Floeter SR, Monteiro-Netto C, Luiz OJ, Ferreira CEL. 2014. Local ecological knowledge and scientific data reveal overexploitation by multigear artisanal fisheries in the Southwestern Atlantic. PLoS ONE 9.

Boaden AE, Kingsford MJ. 2015. Predators drive community structure in coral reef fish assemblages. Ecosphere 6: 1–33.

Bozec YM, O’Farrell S, Bruggemann JH, Luckhurst BE, Mumby PJ. 2016. Tradeoffs between fisheries harvest and the resilience of coral reefs. Proceedings of the National Academy of Sciences of the United States of America 113: 4536–4541.

Burke L, Reytar K, Spalding M, Perry A. 2011. Reefs At Risk Revisited Burkepile DE, Hay ME. 2008. Herbivore species richness and feeding complementarity affect community structure and function on a coral reef. Proceedings of the National Academy of Sciences 105: 16201–16206.

Burkepile DE, Hay ME. 2010. Impact of herbivore identity on algal succession and coral growth on a Caribbean reef. PLoS ONE 5.

Campbell SJ, Edgar GJ, Stuart-Smith RD, Soler G, Bates AE. 2018. Fishing-gear restrictions and biomass gains for coral reef fishes in marine protected areas. Conservation Biology 32: 401–410.

Campbell SJ, Darling ES, Pardede S, Ahmadia G, Mangubhai S, Amkieltiela, Estradivari, Maire E. 2020. Fishing restrictions and remoteness deliver conservation outcomes for Indonesia’s coral reef fisheries. Conservation Letters 13: 1–9.

Cheal AJ, MacNeil MA, Cripps E, Emslie MJ, Jonker M, Schaffelke B, Sweatman H. 2010. Coral – macroalgal phase shifts or reef resilience : links with diversity and functional roles of herbivorous fishes on the Great Barrier Reef. Coral Reefs (2010) 29: 1005–1015.

Chung AE, Wedding LM, Green AL, Friedlander AM, Goldberg G, Meadows A, Hixon MA. 2019. Building Coral reef resilience through spatial herbivore management. Frontiers in Marine Science 6: 1–12.

Cinner JE, Zamborain-Mason J, Gurney GG, Graham NAJ, MacNeil MA, Hoey AS, Mora C, Villéger S, Maire E, McClanahan TR, et al. 2020. Meeting fisheries, ecosystem function, and biodiversity goals in a human-dominated world. Science 368: 307–311.

Clua E, Legendre P. 2008. Shifting dominance among Scarid species on reefs representing a gradient of fishing pressure. Aquatic Living Resources 21: 339–348.

Cure K, Currey-Randall L, Galaiduk R, Radford B, Wakeford M, Heyward A. 2021. Depth gradients in abundance and functional roles suggest limited depth refuges for herbivorous fishes. Coral Reefs 40: 365–379.

Ebrahim A, Martin TSH, Mumby PJ, Olds AD, Tibbetts IR. 2020. Differences in diet and foraging behaviour of commercially important rabbitfish species on coral reefs in the Indian Ocean. Coral Reefs 39: 977–988.

Edwards CB, Friedlander AM, Green AG, Hardt MJ, Sala E, Sweatman HP, Williams ID, Zgliczynski B, Sandin SA, Smith JE. 2014. Global assessment of the status of coral reef herbivorous fishes: Evidence for fishing effects. Proceedings of the Royal Society B: Biological Sciences 281: 7–11.

Fine M, Gildor H, Genin A. 2013. A coral reef refuge in the Red Sea. Global Change Biology 19: 3640– 3647.

Fisher R, O’Leary RA, Low-Choy S, Mengersen K, Knowlton N, Brainard RE, Caley MJ. 2015. Species Richness on Coral Reefs and the Pursuit of Convergent Global Estimates. Current Biology 25: 500– 505.

Floeter SR, Halpern BS, Ferreira CEL. 2006. Effects of fishing and protection on Brazilian reef fishes. Biological Conservation 128: 391–402.

Foo SA, Walsh WJ, Lecky J, Marcoux S, Asner GP. 2021. Impacts of pollution, fishing pressure, and reef rugosity on resource fish biomass in West Hawaii. Ecological Applications 31: 1–15.

Ford AK, Bejarano S, Marshell A, Mumby PJ. 2016. Linking the biology and ecology of key herbivorous unicornfish to fisheries management in the Pacific. Aquatic Conservation: Marine and Freshwater Ecosystems 26: 790–805.

Friedlander AM, Brown EK, Jokiel PL, Smith WR, Rodgers KS. 2003. Effects of habitat, wave exposure, and marine protected area status on coral reef fish assemblages in the Hawaiian archipelago. Coral Reefs 22: 291–305.

Frisch AJ, Cole AJ, Hobbs JPA, Rizzari JR, Munkres KP. 2012. Effects of Spearfishing on Reef Fish Populations in a Multi-Use Conservation Area. PLoS ONE 7.

Galal N. 1999. Studies on the coastal ecology and management of the Nabq Protected Area, South Sinai, Egypt (BL). University of York.

Galal N, Ormond RFG, Hassan O. 2002. Effect of a network of no-take reserves in increasing catch per unit effort and stocks of exploited reef fish at Nabq, South Sinai, Egypt. Marine Freshwater Research 53: 199–205.

Gill DA, Mascia MB, Ahmadia GN, Glew L, Lester SE, Barnes M, Craigie I, Darling ES, Free CM, Geldmann J, et al. 2017. Capacity shortfalls hinder the performance of marine protected areas globally. Nature 543: 665–669.

Golani D, Fricke R. 2018. Checklist of the Red Sea Fishes with delineation of the Gulf of Suez, Gulf of Aqaba, endemism and Lessepsian migrants.

Graham NAJ, Nash KL. 2013. The importance of structural complexity in coral reef ecosystems. Coral Reefs 2012 32:2 32: 315–326.

Green AL, Bellwood DR. 2009. Monitoring functional groups of herbivorous reef fishes as indicators of coral reef resilience : a practical guide for coral reef managers in the Asia Pacific region. No. 7.

Hasler H, Ott JA. 2008. Diving down the reefs? Intensive diving tourism threatens the reefs of the northern Red Sea. Marine Pollution Bulletin 56: 1788–1794.

Hawkins JP, Roberts CM. 2004. Effects of fishing on sex-changing Caribbean parrotfishes. Biological Conservation 115: 213–226.

Heenan A, Williams ID. 2013. Monitoring herbivorous fishes as indicators of coral reef resilience in American Samoa. PLoS ONE 8.

Helyer J, Samhouri JF. 2017. Fishing and environmental influences on estimates of unfished herbivorous fish biomass across the Hawaiian Archipelago. Marine Ecology Progress Series 575: 1–15.

Hernández-Landa RC, Aguilar-Perera A. 2019. Structure and composition of surgeonfish (Acanthuridae) and parrotfish (Labridae: Scarinae) assemblages in the south of the Parque Nacional Arrecife Alacranes, southern Gulf of Mexico. Marine Biodiversity 49: 647–662.

Hixon MA. 2015. Predation: Piscivory and the ecology of coral reef fishes. Ecology of Fishes on Coral Reefs: 41–52.

Hothorn T, Bretz F, Westfall P. 2008. Simultaneous Inference in General Parametric Models. Biometrical Journal 50: 346–363.

Hughes TP, Rodrigues MJ, Bellwood DR, Ceccarelli D, Hoegh-Guldberg O, McCook L, Moltschaniwskyj N, Pratchett MS, Steneck RS, Willis B. 2007. Phase Shifts, Herbivory, and the Resilience of Coral Reefs to Climate Change. Current Biology 17: 360–365.

Humphries AT, McClanahan TR, McQuaid CD. 2020. Algal turf consumption by sea urchins and fishes is mediated by fisheries management on coral reefs in Kenya. Coral Reefs 39: 1137–1146.

Kattan A, Coker DJ, Berumen ML. 2017. Reef fish communities in the central Red Sea show evidence of asymmetrical fishing pressure. Marine Biodiversity 47: 1227–1238.

Khalil MT, Cochran JEM, Berumen ML. 2013. The abundance of herbivorous fish on an inshore Red Sea reef following a mass coral bleaching event. Environmental Biology of Fishes 96: 1065–1072.

Khalil MT, Bouwmeester J, Berumen ML. 2017. Spatial variation in coral reef fish and benthic communities in the central Saudi Arabian Red Sea. PeerJ 2017: 1–23.

Mabrouk A. 2015. THE ROLE OF MARINE PROTECTED AREAS IN MAINTAINING SUSTAINABLE FISHERIES IN THE EGYPTIAN GULF OF AQABA, RED SEA.

Marshell A, Mumby PJ. 2012. Revisiting the functional roles of the surgeonfish Acanthurus nigrofuscus and Ctenochaetus striatus. Coral Reefs 2012 31:4 31: 1093–1101.

Marshell A, Mumby PJ. 2015. The role of surgeonfish (Acanthuridae) in maintaining algal turf biomass on coral reefs. Journal of Experimental Marine Biology and Ecology 473: 152–160.

Maynard JA, McKagan S, Raymundo L, Johnson S, Ahmadia GN, Johnston L, Houk P, Williams GJ, Kendall M, Heron SF, et al. 2015. Assessing relative resilience potential of coral reefs to inform management. Biological Conservation 192: 109–119.

McClanahan TR, Muthiga NA. 2020. Change in fish and benthic communities in Belizean patch reefs in and outside of a marine reserve, across a parrotfish capture ban. Marine Ecology Progress Series 645: 25–40.

McClanahan TR, Muthiga NA, Kamukuru AT, Machano H, Kiambo RW. 1999. The effects of marine parks and fishing on coral reefs of northern Tanzania. Biological Conservation 89: 161–182.

McClanahan TR, Graham NAJ, Macneil MA, Cinner JE. 2015. Biomass-based targets and the management of multispecies coral reef fisheries. Conservation Biology 29: 409–417.

McClanahan TR, Schroeder RE, Friedlander AM, Vigliola L, Wantiez L, Caselle JE, Graham NAJ, Wilson S, Edgar GJ, Stuart-Smith RD, et al. 2019. Global baselines and benchmarks for fish biomass: Comparing remote reefs and fisheries closures. Marine Ecology Progress Series 612: 167–192.

McClanahan TR, Friedlander AM, Graham NAJ, Chabanet P, Bruggemann JH. 2021. Variability in coral reef fish baseline and benchmark biomass in the central and western Indian Ocean provinces. Aquatic Conservation: Marine and Freshwater Ecosystems 31: 28–42.

Mumby PJ, Steneck RS, Adjeroud M, Arnold SN. 2016. High resilience masks underlying sensitivity to algal phase shifts of Pacific coral reefs. Oikos 125: 644–655.

Naumann MS, Bednarz VN, Ferse SCA, Niggl W, Wild C. 2015. Monitoring of coastal coral reefs near Dahab (Gulf of Aqaba, Red Sea) indicates local eutrophication as potential cause for change in benthic communities. Environmental Monitoring and Assessment 187: 1–14.

Olds AD, Albert S, Maxwell PS, Pitt KA, Connolly RM. 2013. Mangrove-reef connectivity promotes the effectiveness of marine reserves across the western Pacific. Global Ecology and Biogeography 22: 1040–1049.

Osman EO, Smith DJ, Ziegler M, Kürten B, Conrad C, El-Haddad KM, Voolstra CR, Suggett DJ. 2018. Thermal refugia against coral bleaching throughout the northern Red Sea. Global Change Biology 24: e474–e484.

Pearson MP, Shehata AI. 1998. Protectorates management for conservation and development in the Arab Republic of Egypt. Parks 8: 29–35.

Poonian C. 2020. Coral reef fisheries of the Mzeina Bedu in South Sinai.

Pratchett MS, Hoey AS, Wilson SK. 2014. Reef degradation and the loss of critical ecosystem goods and services provided by coral reef fishes. Current Opinion in Environmental Sustainability 7: 37– 43.

Price NW, Chen K, Chen CA, Wen CK. 2021. Scraping and grazing herbivorous/detritivorous sh display opposite feeding behaviours under different protection regulations. : 1–22.

Reverter M, Jackson M, Daraghmeh N, von Mach C, Milton N. 2020. 11-yr of coral community dynamics in reefs around Dahab (Gulf of Aqaba, Red Sea): the collapse of urchins and rise of macroalgae and cyanobacterial mats. Coral Reefs 39: 1605–1618.

Robinson JPW, Williams ID, Edwards AM, McPherson J, Yeager L, Vigliola L, Brainard RE, Baum JK. 2017. Fishing degrades size structure of coral reef fish communities. Global Change Biology 23: 1009–1022.

Robinson JPW, McDevitt-Irwin JM, Dajka JC, Hadj-Hammou J, Howlett S, Graba-Landry A, Hoey AS, Nash KL, Wilson SK, Graham NAJ. 2020. Habitat and fishing control grazing potential on coral reefs. Functional Ecology 34: 240–251.

Roff G, Bejarano S, Priest M, Marshell A, Chollett I, Steneck RS, Doropoulos C, Golbuu Y, Mumby PJ. 2019. Seascapes as drivers of herbivore assemblages in coral reef ecosystems. Ecological Monographs 89.

Rolim FA, Langlois T, Rodrigues PFC, Bond T, Motta FS, Neves LM, Gadig OBF. 2018. Network of small no-take marine reserves reveal greater abundance and body size of fisheries target species. bioRxiv: 1–22.

Russ GR, Questel S-LA, Rizzari JR, Alcala AC. 2015. The parrotfish–coral relationship: refuting the ubiquity of a prevailing paradigm. Marine Biology 2015 162:10 162: 2029–2045.

Russ GR, Payne CS, Bergseth BJ, Rizzari JR, Abesamis RA, Alcala AC. 2018. Decadal-scale response of detritivorous surgeonfishes (family Acanthuridae) to no-take marine reserve protection and changes in benthic habitat. Journal of Fish Biology 93: 887–900.

Samy-kamal M. 2015. Status of fisheries in Egypt : reflections on past trends and management challenges. Reviews in Fish Biology and Fisheries 25: 631–649.

Shantz AA, Ladd MC, Burkepile DE. 2020. Overfishing and the ecological impacts of extirpating large parrotfish from Caribbean coral reefs. Ecological Monographs 90: 1–17.

Shpigel M, Fishelson L. 1989. Food habits and prey selection of three species of groupers from the genus Cephalopholis (Serranidae: Teleostei). Environmental Biology of Fishes 24: 67–73.

Shpigel M, Fishelson L. 1991. Experimental removal of piscivorous groupers of the genus Cephalopholis (Serranidae) from coral habitats in the Gulf of Aqaba (Red-Sea). Environmental Biology of Fishes 31: 131–138.

Spalding M, Burke L, Wood SA, Ashpole J, Hutchison J, zu Ermgassen P. 2017. Mapping the global value and distribution of coral reef tourism. Marine Policy 82: 104–113.

Steneck RS, Mumby PJ, MacDonald C, Rasher DB, Stoyle G. 2018. Attenuating effects of ecosystem management on coral reefs. Science Advances 4: 1–12.

Tilot V, Leujak W, Ormond RFG, Ashworth JA, Mabrouk A. 2008. Monitoring of South Sinai coral reefs: Influence of natural and anthropogenic factors. Aquatic Conservation: Marine and Freshwater Ecosystems 18: 1109–1126.

Topor ZM, Rasher DB, Duffy JE, Brandl SJ. 2019. Marine protected areas enhance coral reef functioning by promoting fish biodiversity. Conservation Letters 12: 1–9.

Vallès H, Oxenford HA. 2014. Parrotfish size: A simple yet useful alternative indicator of fishing effects on Caribbean reefs? PLoS ONE 9.

Weijerman M, Gove JM, Williams ID, Walsh WJ, Minton D, Polovina JJ. 2018. Evaluating management strategies to optimise coral reef ecosystem services. Journal of Applied Ecology 55: 1823–1833.

Wilkinson C. 2008. Status of coral reefs of the world: 2008

Williams ID, White DJ, Sparks RT, Lino KC, Zamzow JP, Kelly ELA, Ramey HL. 2016. Responses of Herbivorous Fishes and Benthos to 6 Years of Protection at the Kahekili Herbivore Fisheries Management Area, Maui. PLOS ONE 11: e0159100.

Williams ID, Kindinger TL, Couch CS, Walsh WJ, Minton D, Oliver TA. 2019. Can Herbivore Management Increase the Persistence of Indo-Pacific Coral Reefs? Frontiers in Marine Science 6: 1–15.

Wilson SK, Fisher R, Pratchett MS, Graham NAJ, Dulvy NK, Turner RA, Cakacaka A, Polunin NVC. 2010. Habitat degradation and fishing effects on the size structure of coral reef fish communities. Ecological Applications 20: 442–451.

Zuur AF, Ieno EN, Walker N, Saveliev AA, Smith GM. 2009. Mixed effects models and extensions in ecology with R. Springer New York: New York, NY.

