## Supplementary figures and tables for "Herbivorous fish of Sinai marine protected areas (Gulf of Aqaba): Structure baseline for potential thermal stress impact management"

**This file includes:**

I. Supplementary Figures S1 to S5
II. Supplementary Tables S1 to S10

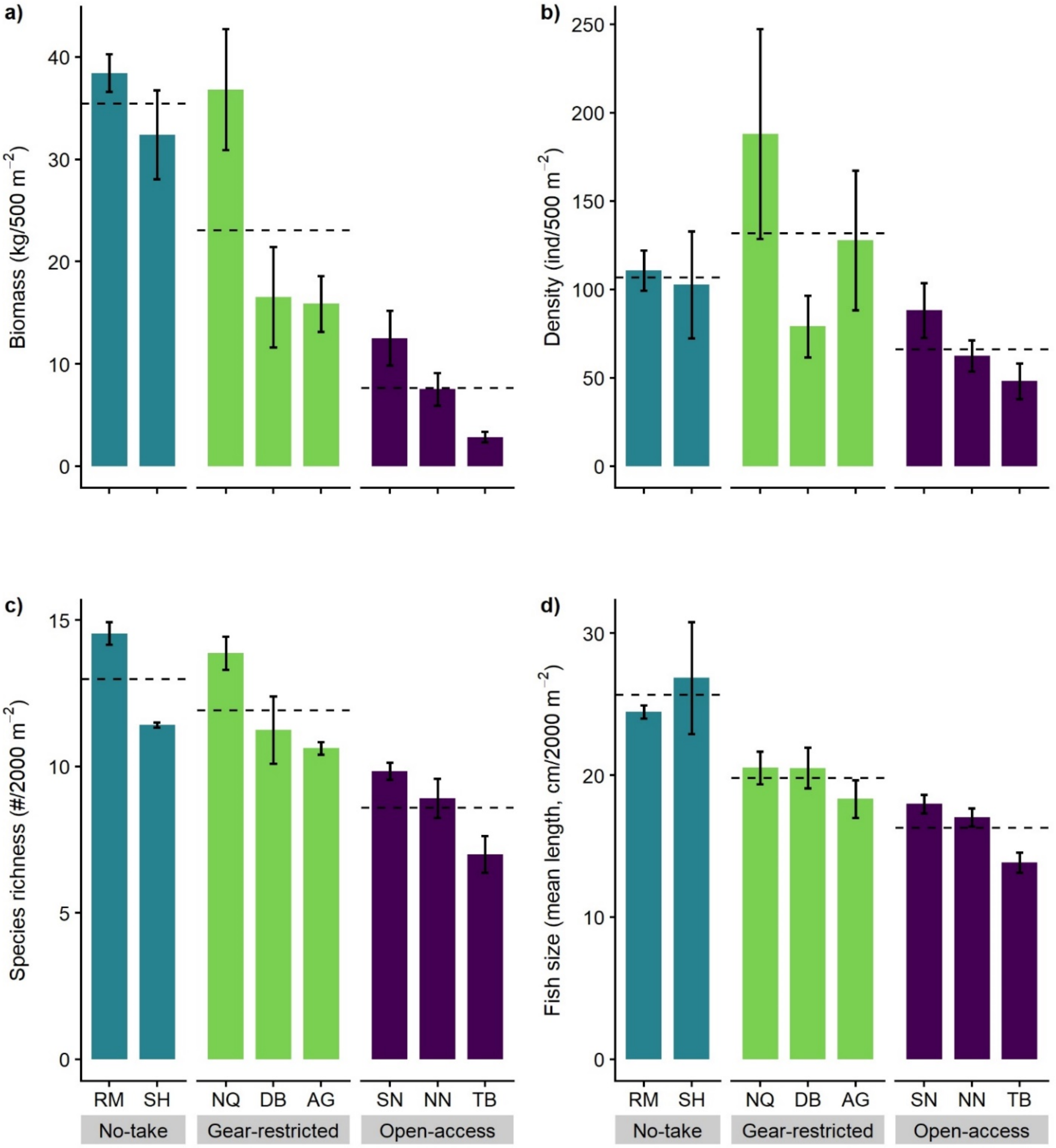


**Figure S1.**

Bar graphs showing the comparison of total herbivorous fish indices between the sampling regions within each fisheries management systems. Fish biomass (a), Fish density (b), Species richness (c) and Fish size (d). Dashed horizontal line represent the overall mean value for each fish index combined within each management systems.

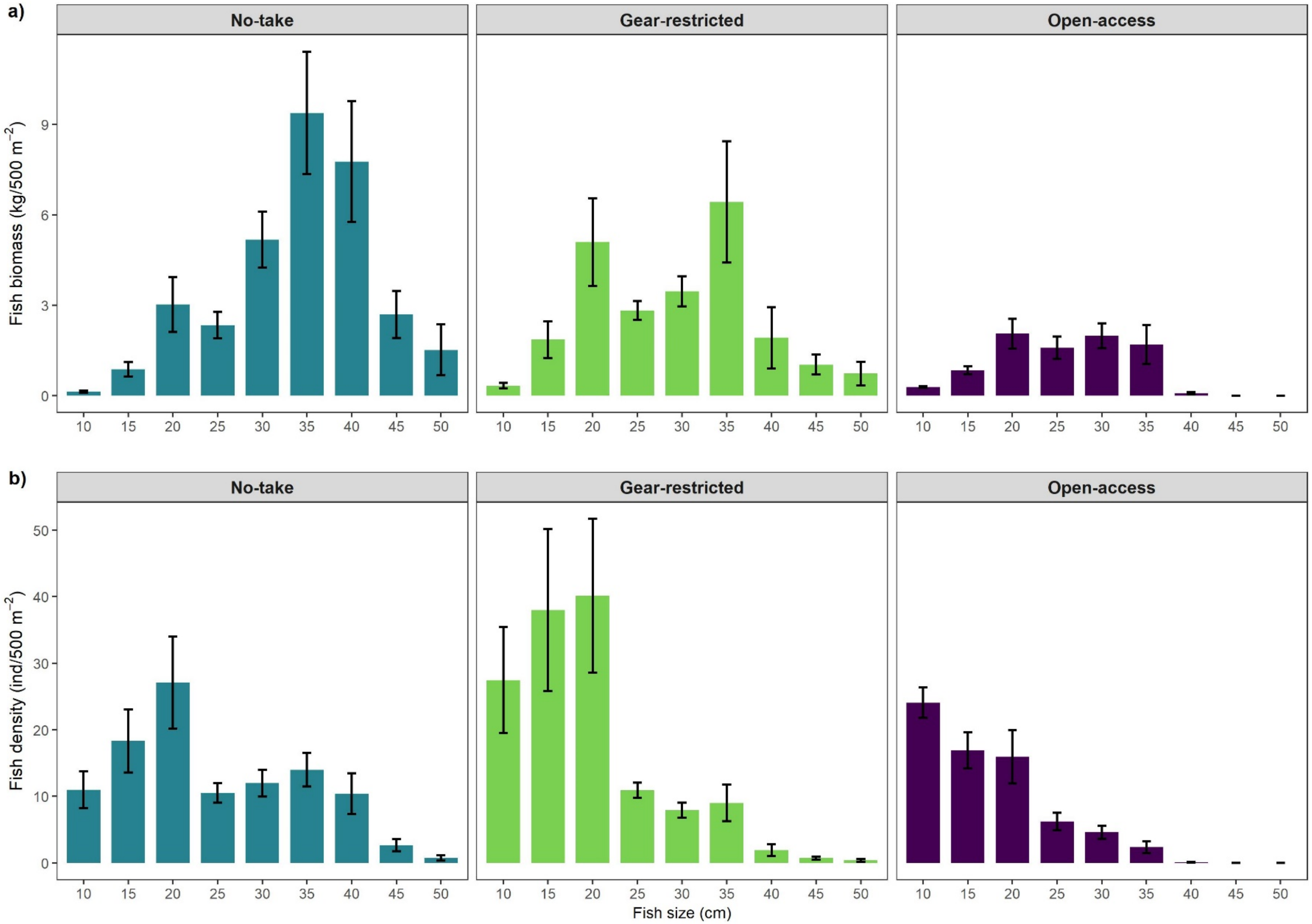


**Figure S2.**

Mean biomass (kg 500$^{-2}$ ± SE) (a) and mean density (individual 500$^{-2}$ ± SE) (b) for each size-class (total length; cm) of total herbivorous reef fishes in each fisheries management systems.

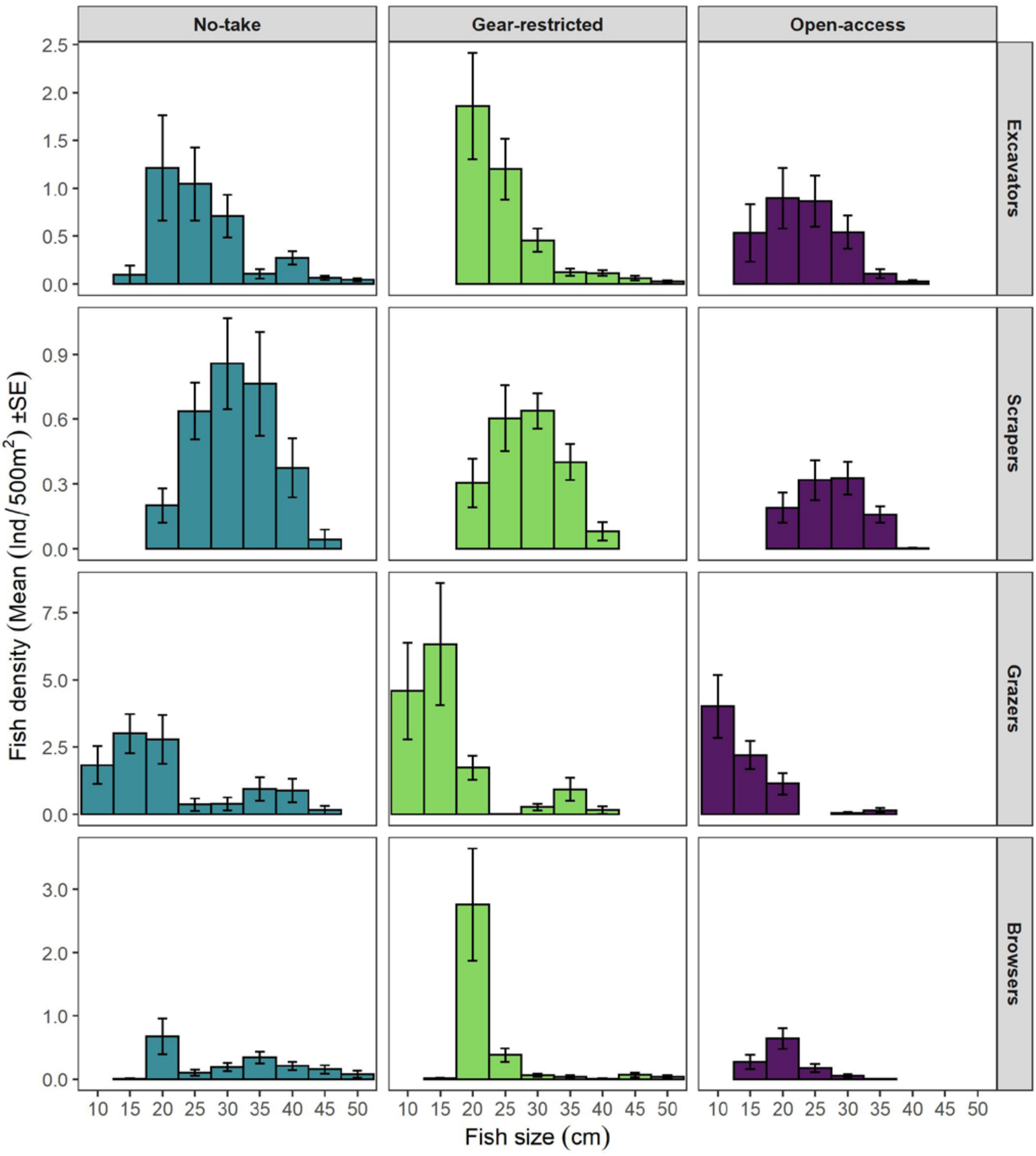


**Figure S3.**

Mean density (individual $500^{-2} \pm$ SE) for each size-class (total length; cm) of herbivorous fishes divided by (functional feeding group) in each fisheries management systems.

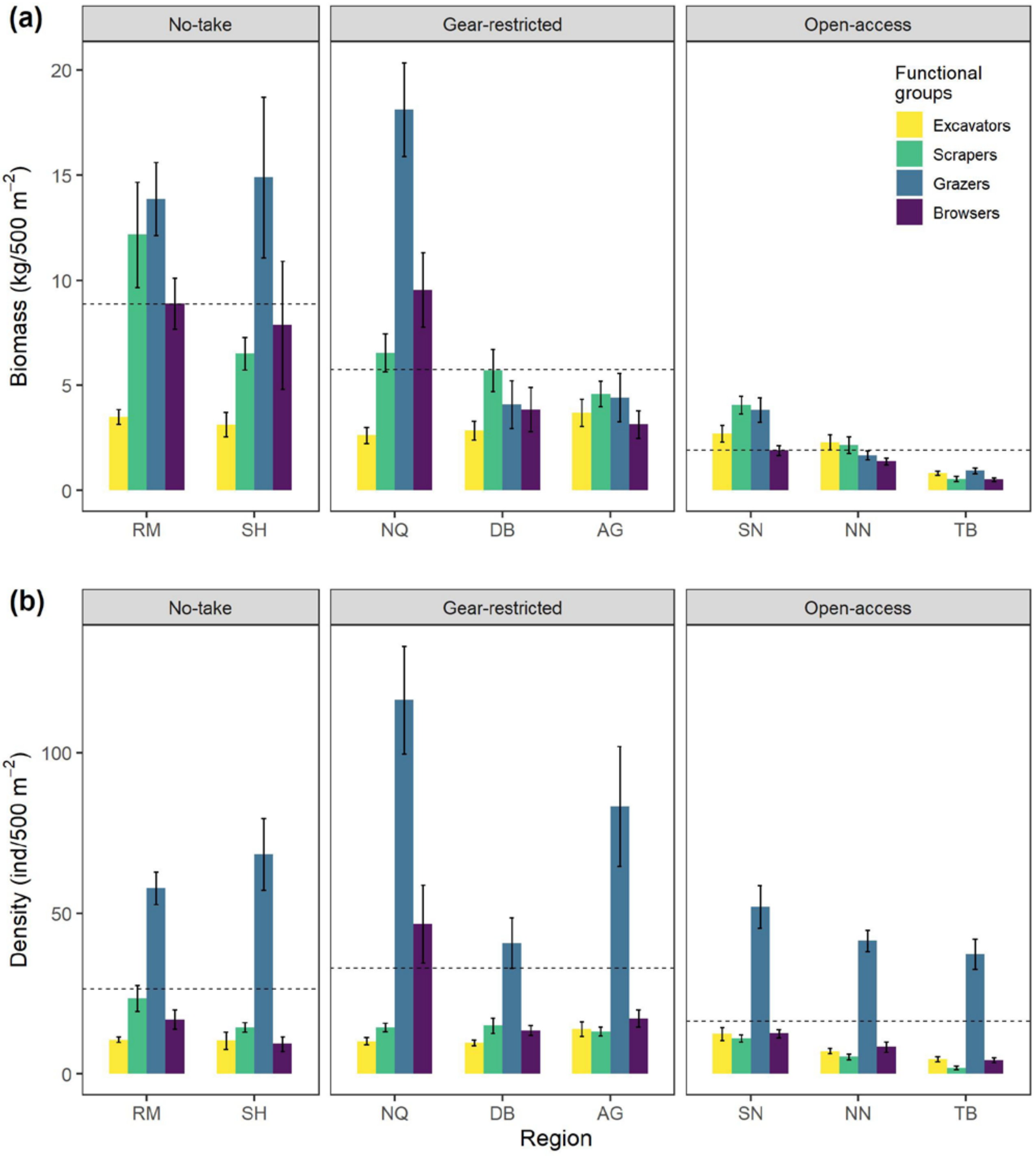


**Figure S4.**

Bar plots, showing the comparison of the four functional groups of herbivores between the sampling regions and management systems. Fish biomass (a), and Fish density (b). Dashed horizontal line represent the overall mean value for each fish metric combined across all regions within each management system.

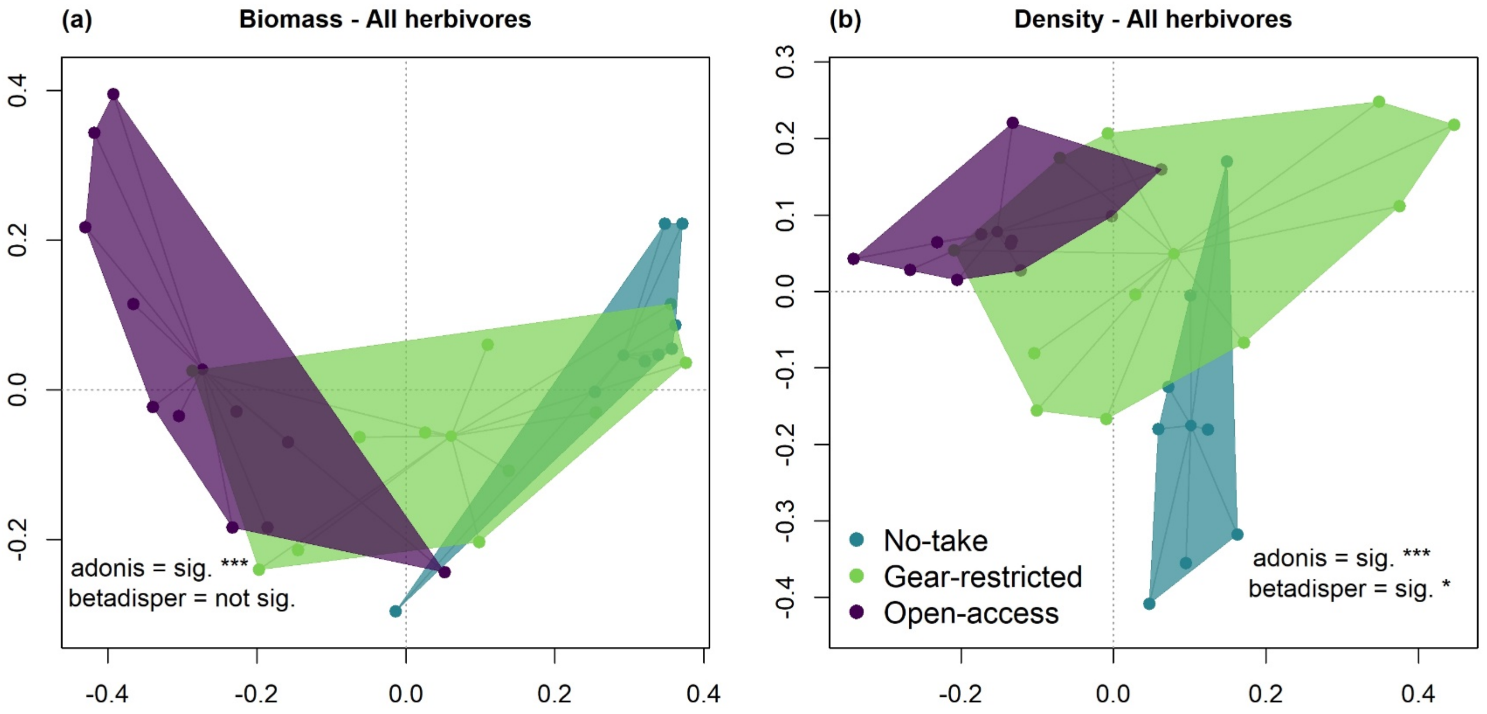


**Figure S5.**

Principal coordinates analysis (PCoA) ordination of distances among the centroids of main effects (fisheries management) based on Bray–Curtis dissimilarities of biomass (a) and density (b) for all herbivores combined. Points represent individual sites (connected to the centroid point in the center), sites are color-coded by fisheries management systems, and shaded polygons indicate boundaries of observed community structure. Significance level of ADONIS and PERMDISP tests are denoted by ***$p$<0.001, **$p$<0.01, *$p$<0.05.

**Table S1.** Details of 30 sites surveyed along the Gulf of Aqaba, Egypt. Eight regions with different type of fisheries management were surveyed, each comprised of 3-5 sites. Sites were assigned a numerical code (1-30) in order of latitude. The coordinates (in decimal degrees format) indicate survey location on reef site.

| Fisheries management | Region | Site ID | Site name | Lat. (N) | Long. (E) |
|---|---|---|---|---|---|
| No-take (NT) | Ras Mohammad (RM) | 1 | Mangrove Channel | 27.724347 | 34.251272 |
| | | 2 | Yollanda Beach | 27.728206 | 34.256730 |
| | | 3 | Old Quay | 27.732875 | 34.242647 |
| | | 4 | Eel Garden | 27.737806 | 34.256753 |
| | | 5 | South Bareika | 27.766686 | 34.220025 |
| | Sharm El-Sheikh (SH) | 6 | Umm el-Sidd | 27.852033 | 34.316664 |
| | | 7 | Sheikh Coast | 27.932728 | 34.369997 |
| | | 8 | North Nasrani | 27.994628 | 34.434483 |
| Gear-restricted (GR) | Nabq (NQ) | 9 | Al-Gharqana | 28.121594 | 34.441964 |
| | | 10 | Maria Schroder | 28.189000 | 34.442275 |
| | | 11 | Ras Tantour | 28.243642 | 34.415519 |
| | | 12 | Al-Sheheira | 28.409100 | 34.451550 |
| | Dahab (DB) | 13 | Three Pools | 28.435117 | 34.456586 |
| | | 14 | Al-Island | 28.477378 | 34.511703 |
| | | 15 | Al-Canyon | 28.556867 | 34.523172 |
| | Abu Galum (AG) | 16 | Al-Dahayla | 28.614869 | 34.559917 |
| | | 17 | Al-Omayed | 28.626903 | 34.575272 |
| | | 18 | Ras Mamlah | 28.731942 | 34.625756 |
| | | 19 | Al-Sokhn | 28.758806 | 34.623367 |
| Open-access (OA) | South Nuweiba (SN) | 20 | Al-Sokhn north | 28.773244 | 34.625372 |
| | | 21 | Wadi Miran | 28.805558 | 34.624556 |
| | | 22 | Hobiq | 28.879167 | 34.645350 |
| | | 23 | Hobiq north | 28.895678 | 34.649092 |
| | | 24 | Al-Mazariq | 28.920800 | 34.644264 |
| | North Nuweiba (NW) | 25 | AL-Tirabin north | 29.063219 | 34.672967 |
| | | 26 | Ras Shattan | 29.126361 | 34.685983 |
| | | 27 | Basata | 29.204842 | 34.735264 |
| | Taba (TB) | 28 | Al-Mahash | 29.325411 | 34.753739 |
| | | 29 | Morgana Beach | 29.359872 | 34.783369 |
| | | 30 | Taba Heights | 29.390611 | 34.810786 |

**Table S2.** List of surveyed fish species, the families they belong to, and the herbivorous functional groups to which they were assigned, with corresponding "a" and "b" values for biomass (kg) calculations, using the equation: $a \times TL^b$ where a and b are constants and total length (cm) is derived from underwater visual census.

| Feeding group | Family | Genus | Species | a | b | Source |
|---|---|---|---|---|---|---|
| Excavators | Scaridae | *Cetoscarus* | *bicolor* | 0.140 | 2.500 | Amin et al. 2019 |
| Excavators | Scaridae | *Chlorurus* | *gibbus* | 0.289 | 2.170 | Amin et al. 2019 |
| Excavators | Scaridae | *Chlorurus* | *sordidus* | 0.109 | 2.480 | Amin et al. 2019 |
| Scrapers | Scaridae | *Hipposcarus* | *harid* | 0.015 | 3.010 | Amin et al. 2019 |
| Scrapers | Scaridae | *Scarus* | *ferrugineus* | 0.028 | 2.880 | Amin et al. 2019 |
| Scrapers | Scaridae | *Scarus* | *frenatus* | 0.129 | 2.816 | Amin et al. 2019 |
| Scrapers | Scaridae | *Scarus* | *fuscopurpureus* | 0.072 | 2.580 | Amin et al. 2019 |
| Scrapers | Scaridae | *Scarus* | *ghobban* | 0.035 | 2.770 | Amin et al. 2019 |
| Scrapers | Scaridae | *Scarus* | *niger* | 0.041 | 2.750 | Amin et al. 2019 |
| Scrapers | Scaridae | *Scarus* | *psittacus* | 0.200 | 2.212 | Froese & Pauly 2019 |
| Grazers | Acanthuridae | *Acanthurus* | *nigrofuscus* | 0.025 | 2.960 | Froese & Pauly 2019 |
| Grazers | Acanthuridae | *Acanthurus* | *sohal* | 0.024 | 2.960 | Froese & Pauly 2019 |
| Grazers | Acanthuridae | *Ctenochaetus* | *striatus* | 0.074 | 2.517 | Afeworki et al. 2013 |
| Grazers | Acanthuridae | *Zebrasoma* | *desjardinii* | 0.023 | 2.960 | Froese & Pauly 2019 |
| Grazers | Acanthuridae | *Zebrasoma* | *xanthurum* | 0.023 | 2.960 | Froese & Pauly 2019 |
| Grazers | Siganidae | *Siganus* | *stellatus* | 0.017 | 2.980 | Froese & Pauly 2019 |
| Browsers | Scaridae | *Calotomus* | *viridescens* | 0.035 | 2.810 | Amin et al. 2019 |
| Browsers | Siganidae | *Siganus* | *luridus* | 0.013 | 3.165 | Abdulghani 2018 |
| Browsers | Siganidae | *Siganus* | *rivulatus* | 0.027 | 2.987 | Gabr et al. 2018 |
| Browsers | Siganidae | *Siganus* | *argenteus* | 0.015 | 2.980 | Froese & Pauly 2019 |
| Browsers | Acanthuridae | *Naso* | *elegans* | 0.022 | 2.970 | Froese & Pauly 2019 |
| Browsers | Acanthuridae | *Naso* | *unicornis* | 0.023 | 2.950 | Froese & Pauly 2019 |
| Browsers | Kyphosidae | *Kyphosus* | *cinerascens* | 0.019 | 3.030 | Froese & Pauly 2019 |
| Browsers | Kyphosidae | *Kyphosus* | *vaigiensis* | 0.017 | 3.000 | Froese & Pauly 2019 |

**Table references:**

Abdulghani, A. A. H. (2018). Biological and ecological processes during the establishment of a marine invasion: the Siganus rabbitfishes from the Red Sea to the coastal areas of Cyrenaica, Libya. PhD Thesis, University of Salford.

Amin, M. A., El-Ganainy, A. A., & Sabrah, M. M. (2019). Length-Weight Relationships of Thirteen Species of Parrotfish (Family Scaridae) inhabiting the Egyptian coasts of the Red Sea. Egyptian Journal of Aquatic Biology and Fisheries, 23(5 (Special Issue)), 357-366.

Gabr MH, Bakaili AS, Mal AO (2018) Growth, mortality and yield per recruit of the rabbit fish Siganus rivulatus (Forsskål 1775) in the red sea coast of Jeddah, Saudi Arabia. International Journal of Fisheries and Aquatic Studies 6:87–96

**Table S3.** Characteristics of herbivore feeding groups. Each herbivorous fish species was assigned one of four feeding groups, defined following established classifications (Green & Bellwood 2009, Lundberg & Lipkin 1979, Ateweberhan et al. 2006, Afeworki et al. 2013, Ebrahim et al. 2020)

| **Feeding group** | **Definition** | **Family** | **Species (n)** |
|---|---|---|---|
| Excavators | Herbivore species that feed on algal turf & associated detritus by excavating surfaces of the reef substratum. | Scaridae | *Cetoscarus bicolor, Chlorurus gibbus, C. sordidus* (3) |
| Scrapers | Herbivore species that feed on algal turf & associated detritus by scraping surfaces of the reef substratum. | Scaridae | *Hipposcarus harid, Scarus ferrugineus, S. frenatus, S. fuscopurpureus, S. ghobban, S. niger, S. psittacus* (7) |
| Grazers | Herbivore species that feed intensely on algal turf & associated detritus without scraping or excavating the reef substratum. | Acanthuridae, Siganidae | *Acanthurus nigrofuscus, A. sohal, Ctenochaetus striatus, Zebrasoma desjardinii, Z. xanthurum, Siganus stellatus* (6) |
| Browsers | Herbivore species that feed on macroalgae & associated epiphytic material. | Scaridae, Siganidae, Acanthuridae, Kyphosidae | *Calotomus viridescens, Siganus luridus, S. rivulatus, S. argenteus Naso unicornis, N. elegans, Kyphosus cinerascens, K. vaigiensis* (8) |

**Table S4.** Functional group, family, and herbivorous fish species observed in the study area. Abundance and biomass are percentages of the grand total. *Freq*. Frequency of occurrence (total number of transects=120). Species are ordered by index of relative dominance (IRD) = (frequency of occurrence × percent biomass) × 100

| **Functional group** | **Family** | **Species** | **Abundance** | **Biomass** | ***Freq*.** | **IRD** |
|---|---|---|---|---|---|---|
| Grazers | Acanthuridae | *Acanthurus sohal* | 6.92 | 25.61 | 55 | 1408.42 |
| Excavators | Scaridae | *Chlorurus sordidus* | 9.06 | 10.23 | 97.5 | 997.37 |
| Scrapers | Scaridae | *Scarus niger* | 5.5 | 7.84 | 96.67 | 757.77 |
| Browsers | Siganidae | *Siganus luridus* | 8.64 | 6.58 | 84.17 | 553.44 |
| Scrapers | Scaridae | *Hipposcarus harid* | 3.08 | 7.58 | 61.67 | 467.2 |
| Grazers | Acanthuridae | *Ctenochaetus striatus* | 13.39 | 4.46 | 98.33 | 438.53 |
| Scrapers | Scaridae | *Scarus ferrugineus* | 2.5 | 5.37 | 79.17 | 424.96 |
| Grazers | Acanthuridae | *Acanthurus nigrofuscus* | 32 | 3.92 | 98.33 | 385.15 |
| Grazers | Acanthuridae | *Zebrasoma xanthurum* | 5.56 | 2.12 | 80 | 169.35 |
| Browsers | Acanthuridae | *Naso unicornis* | 1.05 | 7.46 | 21.67 | 161.7 |
| Scrapers | Scaridae | *Scarus frenatus* | 0.58 | 3.86 | 35.83 | 138.39 |
| Browsers | Acanthuridae | *Naso elegans* | 1.04 | 3.31 | 33.33 | 110.31 |
| Grazers | Acanthuridae | *Zebrasoma desjardinii* | 3.07 | 1.65 | 66.67 | 109.99 |
| Browsers | Scaridae | *Calotomus viridescens* [a] | 1.55 | 1.57 | 65.83 | 103.27 |
| Excavators | Scaridae | *Chlorurus gibbus* [a] | 0.64 | 1.92 | 40 | 76.75 |
| Scrapers | Scaridae | *Scarus fuscopurpureus* | 0.56 | 0.99 | 25 | 24.75 |
| Browsers | Siganidae | *Siganus rivulatus* | 2.59 | 1.09 | 20.83 | 22.78 |
| Excavators | Scaridae | *Cetoscarus bicolor* | 0.17 | 0.87 | 14.17 | 12.32 |
| Browsers | Kyphosidae | *Kyphosus vaigiensis* | 0.29 | 1.17 | 8.33 | 9.74 |
| Browsers | Siganidae | *Siganus argenteus* | 1.02 | 0.76 | 6.67 | 5.07 |
| Scrapers | Scaridae | *Scarus ghobban* | 0.24 | 0.66 | 7.5 | 4.93 |
| Browsers | Kyphosidae | *Kyphosus cinerascens* | 0.17 | 0.7 | 6.67 | 4.67 |
| Scrapers | Scaridae | *Scarus psittacus* | 0.16 | 0.22 | 10 | 2.18 |
| Grazers | Siganidae | *Siganus stellatus* | 0.22 | 0.08 | 5 | 0.41 |

[a] Endemic species to Red Sea

**Table S5.** Comparisons of fish biomass (kg 500 $m^{-2}$) and density (individual 500 $m^{-2}$) of the studied functional groups (excavators, scrapers, grazers, and browsers) and for all groups combined between the surveyed regions and fisheries management systems. Values are grand means (±SE). Fisheries management systems abbreviations: NT = No-take, GR = Gear-restricted, OA= Open-access. Region abbreviations: RM = Ras Mohammad, SH= Sharm El-Sheikh, NQ= Nabq, DB= Dahab, AG= Abu Galum, SN= South Nuweiba, NN= North Nuweiba, TB= Taba

| | **RM (n= 5)** | **SH (n= 3)** | **Total NT** | **NQ (n= 4)** | **DB (n= 3)** | **AG (n= 4)** | **Total GR** | **SN (n= 5)** | **NN (n= 3)** | **TB (n= 3)** | **Total OA** |
|---|---|---|---|---|---|---|---|---|---|---|---|
| (a) BIOMASS | | | | | | | | | | | |
| Excavators | 3.5 | 3.1 | **3.4** | 2.6 | 2.9 | 3.7 | **3.1** | 2.7 | 2.3 | 0.8 | **2.1** |
| SE | ±0.2 | ±1.0 | **±0.4** | ±0.7 | ±0.9 | ±0.4 | **±0.4** | ±0.8 | ±0.7 | ±0.2 | **±0.4** |
| Scrapers | 12.2 | 6.5 | **10.0** | 6.5 | 5.7 | 4.6 | **5.6** | 4.1 | 2.2 | 0.6 | **2.6** |
| SE | ±2.6 | ±1.6 | **±1.9** | ±1.7 | ±1.2 | ±0.6 | **±0.7** | ±0.6 | ±0.8 | ±0.2 | **±0.6** |
| Grazers | 13.9 | 14.9 | **14.3** | 18.1 | 4.1 | 4.4 | **9.3** | 3.8 | 1.7 | 0.9 | **2.5** |
| SE | ±2.7 | ±5.6 | **±2.4** | ±4.7 | ±2.1 | ±2.3 | **±2.8** | ±1.0 | ±0.5 | ±0.2 | **±0.6** |
| Browsers | 8.9 | 7.9 | **8.5** | 9.5 | 3.9 | 3.1 | **5.7** | 1.9 | 1.4 | 0.5 | **1.4** |
| SE | ±1.5 | ±6.7 | **±2.4** | ±2.5 | ±1.2 | ±0.6 | **±1.3** | ±0.5 | ±0.1 | ±0.1 | **±0.3** |
| All herbivores | 38.4 | 32.4 | **36.2** | 36.8 | 16.5 | 15.9 | **23.7** | 12.5 | 7.5 | 2.9 | **8.5** |
| SE | ±1.8 | ±4.3 | **±2.1** | ±5.9 | ±4.9 | ±2.7 | **±4.0** | ±2.7 | ±1.6 | ±0.5 | **±1.8** |
| (b) DENSITY | | | | | | | | | | | |
| Excavators | 10.7 | 10.4 | **10.6** | 11.1 | 10.6 | 13.3 | **11.7** | 13.5 | 7.0 | 6.6 | **9.9** |
| SE | ±1.0 | ±5.5 | **±1.9** | ±2.2 | ±1.5 | ±2.0 | **±1.1** | ±4.0 | ±0.3 | ±3.5 | **±2.2** |
| Scrapers | 23.6 | 14.5 | **20.2** | 14.3 | 17.3 | 11.7 | **14.2** | 11.6 | 2.9 | 5.8 | **7.6** |
| SE | ±6.3 | ±3.3 | **±4.3** | ±2.2 | ±1.9 | ±1.6 | **±1.2** | ±1.3 | ±1.4 | ±4.0 | **±1.6** |
| Grazers | 57.9 | 68.4 | **61.9** | 107.2 | 43.2 | 90.4 | **83.6** | 52.9 | 40.9 | 38.5 | **47.0** |
| SE | ±8.6 | ±24.6 | **±9.7** | ±33.5 | ±10.7 | ±38.3 | **±18.8** | ±11.3 | ±5.9 | ±9.3 | **±6.0** |
| Browsers | 17.0 | 9.4 | **14.2** | 52.2 | 14.9 | 19.8 | **30.2** | 13.3 | 5.9 | 7.7 | **9.7** |
| SE | ±3.7 | ±4.7 | **±3.0** | ±24.2 | ±1.0 | ±1.4 | **±9.6** | ±1.0 | ±1.3 | ±4.4 | **±1.6** |
| All herbivores | 109.1 | 102.7 | **106.7** | 188.1 | 79.2 | 127.9 | **136.5** | 88.2 | 62.4 | 48.2 | **70.2** |
| SE | ±10.8 | ±30.2 | **±11.9** | ±59.4 | ±17.5 | ±39.5 | **±27.6** | ±15.5 | ±8.7 | ±10.0 | **±9.1** |

**Table S6.** Comparisons of species richness (number 2000 $m^{-2}$) and body-size (length in cm 2000 $m^{-2}$) of the studied functional groups (excavators, scrapers, grazers, and browsers) and for all groups combined between the surveyed regions and fisheries management systems. Values are grand means (±SE). Fisheries management systems abbreviations: NT = No-take, GR = Gear-restricted, OA= Open-access. Region abbreviations: RM = Ras Mohammad, SH= Sharm El-Sheikh, NQ= Nabq, DB= Dahab, AG= Abu Galum, SN= South Nuweiba, NN= North Nuweiba, TB= Taba

| | **RM (n= 5)** | **SH (n= 3)** | **Total NT** | **NQ (n= 4)** | **DB (n= 3)** | **AG (n= 4)** | **Total GR** | **SN (n= 5)** | **NN (n= 3)** | **TB (n= 3)** | **Total OA** |
|---|---|---|---|---|---|---|---|---|---|---|---|
| (a) SPECIES RICHNESS | | | | | | | | | | | |
| Excavators | 3.0 | 2.3 | **2.8** | 2.3 | 2.0 | 2.5 | **2.3** | 1.6 | 1.7 | 1.0 | **1.5** |
| SE | ±0.0 | ±0.3 | **±0.2** | ±0.5 | ±0.6 | ±0.3 | **±0.2** | ±0.2 | ±0.3 | ±0.0 | **±0.2** |
| Scrapers | 5.6 | 3.7 | **4.9** | 6.0 | 4.3 | 4.5 | **5.0** | 3.8 | 3.3 | 2.3 | **3.3** |
| SE | ±0.4 | ±0.3 | **±0.4** | ±0.3 | ±0.3 | ±0.6 | **±0.4** | ±0.2 | ±1.3 | ±0.7 | **±0.4** |
| Grazers | 5.2 | 4.7 | **5.0** | 5.0 | 4.7 | 4.3 | **4.6** | 4.2 | 4.0 | 4.3 | **4.2** |
| SE | ±0.2 | ±0.3 | **±0.2** | ±0.4 | ±0.3 | ±0.5 | **±0.2** | ±0.4 | ±0.0 | ±0.3 | **±0.2** |
| Browsers | 5.2 | 2.7 | **4.3** | 4.3 | 3.3 | 3.8 | **3.8** | 2.4 | 2.7 | 2.0 | **2.4** |
| SE | ±0.5 | ±0.7 | **±0.6** | ±0.3 | ±0.7 | ±0.8 | **±0.3** | ±0.2 | ±0.3 | ±0.0 | **±0.2** |
| All herbivores | 19.0 | 13.3 | **16.9** | 17.5 | 14.3 | 15.0 | **15.7** | 12.0 | 11.7 | 9.7 | **11.3** |
| SE | ±0.7 | ±0.3 | **±1.1** | ±0.9 | ±1.8 | ±1.1 | **±0.8** | ±0.3 | ±1.8 | ±0.3 | **±0.5** |
| (b) BODY-SIZE (cm) | | | | | | | | | | | |
| Excavators | 26.6 | 29.1 | **27.5** | 24.0 | 24.9 | 24.7 | **24.5** | 22.9 | 22.5 | 25.0 | **23.4** |
| SE | ±0.9 | ±2.8 | **±1.2** | ±0.9 | ±0.9 | ±0.8 | **±0.5** | ±0.7 | ±1.0 | ±1.5 | **±0.6** |
| Scrapers | 31.8 | 32.0 | **31.9** | 29.7 | 29.0 | 26.8 | **28.4** | 27.4 | 28.5 | 27.6 | **27.8** |
| SE | ±1.1 | ±2.3 | **±1.0** | ±0.9 | ±1.0 | ±0.9 | **±0.6** | ±0.8 | ±0.5 | ±0.6 | **±0.4** |
| Grazers | 23.0 | 23.1 | **23.0** | 17.8 | 15.4 | 14.2 | **15.8** | 14.1 | 14.2 | 14.4 | **14.2** |
| SE | ±3.9 | ±4.0 | **±2.7** | ±1.2 | ±1.7 | ±0.4 | **±0.8** | ±0.6 | ±1.1 | ±0.3 | **±0.4** |
| Browsers | 30.2 | 32.4 | **31.0** | 23.6 | 23.3 | 21.4 | **22.7** | 20.2 | 21.1 | 19.6 | **20.3** |
| SE | ±2.2 | ±5.2 | **±2.2** | ±2.0 | ±1.5 | ±0.2 | **±0.8** | ±0.9 | ±1.2 | ±0.5 | **±0.5** |
| All herbivores | 24.5 | 26.9 | **25.4** | 20.5 | 20.5 | 18.3 | **19.7** | 18.0 | 17.0 | 13.9 | **16.6** |
| SE | ±0.5 | ±3.9 | **±1.4** | ±1.2 | ±1.4 | ±1.3 | **±0.7** | ±0.6 | ±0.6 | ±0.7 | **±0.6** |

**Table S7.** Result of the ADONIS and Pairwise comparison tests examining the effect of fisheries management systems on the biomass and density of herbivorous fish communities in reef crest habitat along the Gulf of Aqaba, Egypt. Site × species matrices were calculated using fish biomass and density of 24 herbivorous fish species based on Bray-Curtis distance matrices. Permutation test for homogeneity of multivariate dispersions [a] (PERMDISP). Pairwise ADONIS investigating the compositional differences in the biomass and density of herbivorous fish communities among three fisheries management systems (NT: No-take, GR: Gear-restricted and OA: Open-access). *P-F*: *Pseudo-F*; Bold values: indicate significant effect at $P < 0.05$

| | | | | | Pairwise ADONIS | | | | | | | | |
|---|---|---|---|---|---|---|---|---|---|---|---|---|---|
| | ADONIS | | | PERMDISP | NT vs OA | | | NT vs GR | | | GR vs OA | | |
| | *P-F* | $R^2$ | *P(perm)* | *P(perm)* | F model | $R^2$ | *p.adj* | F model | $R^2$ | *p.adj* | F model | $R^2$ | *p.adj* |
| (a) Biomass | | | | | | | | | | | | | |
| All herbivores | 7.749 | 0.32 | 0.001 | 0.45 | 11.424 | 0.40 | 0.003 | 3.121 | 0.16 | 0.048 | 5.161 | 0.21 | 0.006 |
| Excavators | 3.896 | 0.21 | 0.01 | 0.32 | 5.287 | 0.24 | 0.03 | 0.765 | 0.04 | 1 | 3.720 | 0.16 | 0.11 |
| Scrapers | 4.611 | 0.21 | 0.001 | 0.06 | 4.785 | 0.22 | 0.003 | 1.648 | 0.09 | 0.53 | 3.755 | 0.16 | 0.006 |
| Grazers | 6.402 | 0.28 | 0.001 | 0.008 | 11.559 | 0.41 | 0.003 | 2.910 | 0.15 | 0.12 | 3.350 | 0.14 | 0.03 |
| Browsers | 7.517 | 0.35 | 0.001 | 0.04 | 11.915 | 0.41 | 0.003 | 3.405 | 0.17 | 0.02 | 7.163 | 0.26 | 0.003 |
| (b) Density | | | | | | | | | | | | | |
| All herbivores | 4.776 | 0.25 | 0.001 | 0.02 | 7.487 | 0.31 | 0.003 | 3.057 | 0.15 | 0.02 | 3.952 | 0.17 | 0.01 |
| Excavators | 1.634 | 0.10 | 0.2 | 0.69 | 1.292 | 0.07 | 0.765 | 0.725 | 0.04 | 1 | 2.463 | 0.11 | 0.22 |
| Scrapers | 4.332 | 0.20 | 0.001 | 0.08 | 4.437 | 0.21 | 0.003 | 1.029 | 0.06 | 1 | 3.918 | 0.16 | 0.01 |
| Grazers | 4.103 | 0.23 | 0.002 | 0.004 | 7.706 | 0.31 | 0.003 | 2.814 | 0.14 | 0.08 | 3.114 | 0.14 | 0.045 |
| Browsers | 5.026 | 0.27 | 0.001 | 0.045 | 7.397 | 0.30 | 0.003 | 4.019 | 0.19 | 0.009 | 3.911 | 0.16 | 0.02 |

[a] The multivariate dispersions (number of permutations = 999) indicate that the dispersion of the distances in herbivorous fish communities among the three fisheries management systems is significantly heterogeneous.

**Table S8.** Result of PERMANOVA and PERMDISP comparing the variability in herbivorous fish density and biomass among fisheries management and among the spatial factors (region and site) based on the Bray-Curtis similarity index for square-root transformed fish density and biomass data of 24 species from the reef crest zone along the Gulf of Aqaba, Egypt. *P*-values were obtained using 9999 permutations under a reduced model. Each term is identified as contributing either a fixed or random component to the overall model; df: degree of freedom; ECV: Estimate Components of Variation; Sqrt: square root of ECV. Bold values: indicate significant results at $P < 0.05$

| **Source** | **PERMANOVA** | | | | | | **PERMDISP** | |
|---|---|---|---|---|---|---|---|---|
| | df | *Pseudo F* | *P(perm)* | Component | ECV | Sqrt | *F* | *P(perm)* |
| (a) Biomass | | | | | | | | |
| Management | 2 | 4.1971 | 0.0018 | Fixed | 415.04 | 20.373 | 2.6378 | 0.0901 |
| Region (Management) | 5 | 2.1098 | 0.0002 | Random | 178.41 | 13.357 | 5.5461 | 0.0002 |
| Site (Region (Management)) | 22 | 6.2498 | 0.0001 | Random | 489.21 | 22.118 | 4.9832 | 0.0001 |
| Residual | 90 | | | | 372.74 | 19.307 | | |
| Total | 119 | | | | | | | |
| (b) Density | | | | | | | | |
| Management | 2 | 4.4123 | 0.0003 | Fixed | 261 | 16.156 | 7.0306 | 0.003 |
| Region (Management) | 5 | 1.6081 | 0.0129 | Random | 75.534 | 8.691 | 6.6802 | 0.0001 |
| Site (Region (Management)) | 22 | 7.7759 | 0.0001 | Random | 392.1 | 19.802 | 6.7281 | 0.0001 |
| Residual | 90 | | | | 231.47 | 15.214 | | |
| Total | 119 | | | | | | | |

**Table S9.** Results from similarity of percentages analysis (SIMPER) of discriminating species contributing to 70% of the variation in herbivorous fish community structure between fisheries management groups. Mean fish biomass (kg 500m$^2$) as well as the cumulative contributions (%) of each species to overall community dissimilarity between management groups are given

| Species | Functional group | Mean biomass | Mean biomass | Contribution % | Cumulative % |
|---|---|---|---|---|---|
| | | No-take | Open-access | | |
| *Acanthurus sohal* | Grazers | 45.48 | 3.16 | 32.6 | 32.6 |
| *Naso unicornis* | Browsers | 15.65 | 0.00 | 11.3 | 43.9 |
| *Hipposcarus harid* | Scrapers | 14.61 | 1.67 | 9.6 | 53.5 |
| *Naso elegans* | Browsers | 8.33 | 0.25 | 5.9 | 59.4 |
| *Scarus ferrugineus* | Scrapers | 7.98 | 1.38 | 5.0 | 64.4 |
| *Scarus niger* | Scrapers | 8.06 | 5.20 | 4.8 | 69.2 |
| *Ctenochaetus striatus* | Grazers | 6.34 | 3.52 | 4.4 | 73.6 |
| | | No-take | Gear-restricted | | |
| *Acanthurus sohal* | Grazers | 45.48 | 23.69 | 29.0 | 29.0 |
| *Naso unicornis* | Browsers | 15.65 | 6.08 | 11.3 | 40.3 |
| *Hipposcarus harid* | Scrapers | 14.61 | 5.43 | 9.2 | 49.5 |
| *Siganus luridus* | Browsers | 3.20 | 9.89 | 5.4 | 54.9 |
| *Naso elegans* | Browsers | 8.33 | 1.43 | 5.4 | 60.3 |
| *Scarus niger* | Scrapers | 8.06 | 7.28 | 4.2 | 64.5 |
| *Scarus frenatus* | Scrapers | 6.01 | 2.70 | 4.1 | 68.6 |
| *Ctenochaetus striatus* | Grazers | 6.34 | 2.30 | 4.1 | 72.7 |
| | | Gear-restricted | Open-access | | |
| *Acanthurus sohal* | Grazers | 23.69 | 3.16 | 22.5 | 22.5 |
| *Siganus luridus* | Browsers | 9.89 | 3.17 | 8.9 | 31.4 |
| *Naso unicornis* | Browsers | 6.08 | 0.00 | 8.9 | 40.3 |
| *Chlorurus sordidus* | Excavators | 9.30 | 7.74 | 7.5 | 47.8 |
| *Scarus niger* | Scrapers | 7.28 | 5.20 | 6.8 | 54.6 |
| *Hipposcarus harid* | Scrapers | 5.43 | 1.67 | 6.1 | 60.7 |
| *Scarus ferrugineus* | Scrapers | 5.38 | 1.38 | 6.1 | 66.8 |
| *Acanthurus nigrofuscus* | Grazers | 6.57 | 1.14 | 6.1 | 72.9 |

**Table S10.** Results from similarity of percentages analysis (SIMPER) of discriminating species contributing to 70% of the variation in herbivorous fish community structure between fisheries management groups. Mean fish density (individual 500m$^2$) as well as the cumulative contributions (%) of each species to overall community dissimilarity between management groups are given

| **Species** | **Functional group** | **Mean density** | **Mean density** | **Contribution %** | **Cumulative %** |
|---|---|---|---|---|---|
| | | No-take | Open-access | | |
| *Acanthurus sohal* | Grazers | 57.25 | 4.64 | 16.0 | 16.0 |
| *Ctenochaetus striatus* | Grazers | 85.37 | 54.00 | 15.9 | 31.9 |
| *Acanthurus nigrofuscus* | Grazers | 71.25 | 96.27 | 14.6 | 46.5 |
| *Hipposcarus harid* | Scrapers | 27.62 | 4.09 | 6.3 | 52.8 |
| *Chlorurus sordidus* | Excavators | 37.62 | 34.27 | 6.2 | 59.0 |
| *Siganus luridus* | Browsers | 19.37 | 24.18 | 6.1 | 65.1 |
| *Zebrasoma desjardinii* | Grazers | 20.75 | 5.36 | 4.6 | 69.7 |
| *Scarus ferrugineus* | Scrapers | 17.37 | 3.36 | 4.1 | 73.8 |
| | | No-take | Gear-restricted | | |
| *Acanthurus nigrofuscus* | Grazers | 71.25 | 215.82 | 26.8 | 26.8 |
| *Ctenochaetus striatus* | Grazers | 85.37 | 36.18 | 12.3 | 39.1 |
| *Acanthurus sohal* | Grazers | 57.25 | 32.45 | 11.0 | 50.1 |
| *Siganus luridus* | Browsers | 19.37 | 60.00 | 8.6 | 58.7 |
| *Zebrasoma xanthurum* | Grazers | 9.37 | 36.00 | 6.5 | 65.2 |
| *Chlorurus sordidus* | Excavators | 37.62 | 41.36 | 4.4 | 69.6 |
| *Hipposcarus harid* | Scrapers | 27.62 | 10.82 | 4.3 | 73.9 |
| | | Gear-restricted | Open-access | | |
| *Acanthurus nigrofuscus* | Grazers | 215.82 | 96.27 | 30.8 | 30.8 |
| *Ctenochaetus striatus* | Grazers | 36.18 | 54.00 | 9.4 | 40.2 |
| *Siganus luridus* | Browsers | 60.00 | 24.18 | 9.4 | 49.6 |
| *Zebrasoma xanthurum* | Grazers | 36.00 | 20.36 | 8.3 | 57.9 |
| *Acanthurus sohal* | Grazers | 32.45 | 4.64 | 7.5 | 65.4 |
| *Chlorurus sordidus* | Excavators | 41.36 | 34.27 | 6.4 | 71.8 |
